# The role of the nuclear pore complex in the stability of disease-related short tandem DNA repeats

**DOI:** 10.64898/2026.01.15.699629

**Authors:** Sandra Ollivaud, Daniele Novarina, Elizabeth C. Riquelme Barrientos, Hinke G. Kazemier, Suzan Gonera, Maxime Kislanski, Liesbeth M. Veenhoff, Michael Chang

**Author notes:** Correspondence to Liesbeth M. Veenhoff; Michael Chang.

## Abstract

Nuclear pore complexes (NPCs) mediate selective transport between the nucleus and the cytoplasm, but also contribute to maintaining genome stability. Mutations in NPC genes cause genome instability and sensitivity to DNA damaging agents, and DNA that is difficult to repair or replicate relocates to NPCs, including expanded CAG repeats, which are associated with several neurological diseases. Here, we show that other disease-related short tandem repeats also relocate to NPCs. Relocation depends on the NPC basket protein Nup1, but is independent of the rest of the basket. Abrogating relocation to the NPC increases rates of repeat contraction, but not expansion. By contrast, deletion of the basket component *NUP60,* which causes mislocalization of all other basket proteins except Nup1, results in greater genome instability without affecting relocation to NPCs. Our results show that NPC association is a general feature of disease-related short tandem repeats, and suggest that relocation to NPCs is separable from the other genome maintenance functions of the NPC basket.

## Introduction

Short tandem repeats (STRs) are present in all domains of life, comprise around 7% of the human genome, and are highly variable between individuals (Gemayel et al., 2010; Shortt et al., 2020). This variability is thought to stem from their ability to contract and expand. STRs form noncanonical secondary structures, such as G-quadruplexes, hairpins, triplexes, and DNA unwinding elements, which can obstruct DNA replication, repair, or transcription in a manner that results in the expansion or contraction of STRs (Khristich and Mirkin, 2020; Murat et al., 2020).

While STRs serve important structural and regulatory roles in the genome, their instability can have profound pathological consequences, particularly when expanded beyond a certain threshold. Expansion of some STRs beyond a certain gene- and context-dependent threshold can cause neurological disorders, such as Huntington’s disease or Friedreich’s ataxia (Khristich and Mirkin, 2020). To date, expansion of 15 sequences are known to cause around 80 repeat expansion disorders (REDs; Rajan-Babu et al., 2024). Notably, this number continues to grow with advances in sequencing technologies allowing detection of expanded STRs (Tanudisastro et al., 2024). Furthermore, many cancers exhibit STR instability near regulatory elements, a phenomenon thought to support aberrant cell proliferation (Erwin et al., 2023). A systematic analysis of STRs may help disentangle general mechanisms of instability from those that are sequence- or structure-specific. Recent attention has turned to the nuclear pore complex (NPC) emerging as a potential modulator of repeat stability.

The NPC is most well known for its role as the selective transport channel between the nucleus and cytoplasm, but it has additional roles in genome organization and stability (Simon et al., 2024). NPCs, broadly conserved across eukaryotes, are large protein complexes composed of around 30 different NPC proteins (nucleoporins or Nups). The nuclear side of the NPC—namely the nuclear basket (a dynamic and flexible structure composed of Nup60, Nup1, Nup2, Mlp1, Mlp2, and Pml39 in the baker’s yeast *Saccharomyces cerevisiae*) and the outer rings (consisting of the Nup84-complex composed of Nup84, Nup85, Nup120, Nup133, Nup145C, Sec13 and Seh1)—has contact with the genome, creating a hub for various processes to take place, including chromatin organization, gene expression and DNA repair (Strambio-De-Castillia et al., 2010; Beck and Hurt, 2017; Simon et al., 2024). Strains lacking outer ring proteins (Nup84, Nup120, Nup133) or basket proteins (Nup60, Mlp1, Mlp2) are sensitive to DNA damaging agents and are synthetic lethal or sick in the absence of double-strand break (DSB) repair proteins (Bennett et al., 2001; Chang et al., 2002; Hanway et al., 2002; Kölling et al., 1993; Palancade et al., 2007). The SUMO protease Ulp1 attaches to the nuclear basket to mediate at least some of these effects as Ulp1 is mislocalized and reduced in abundance in *nup60Δ* and *mlp1Δ mlp2Δ* strains, and overexpression of Ulp1 can suppress some of the defects of these strains (Zhao et al., 2004; Palancade et al., 2007). In mammalian cells, it was shown that Nup153 (homolog of yeast Nup60 and Nup1) has a role in DSB repair and the activation of the G2/M and S phase checkpoints (Matsuoka et al., 2007; Lemaître et al., 2012; Duheron et al., 2017), while TPR (Mlp1 and Mlp2 homolog) is important for preventing transcription-mediated replication stress, like R-loops, and maintaining nuclear organization during embryogenesis (Kosar et al., 2021; Wu et al., 2021). Nup107 (Nup84 homolog) is involved in the DNA damage response, by facilitating nuclear translocation of Apaf-1 to mediate the S phase checkpoint response (Jagot-Lacoussiere et al., 2015), indicating that the genome maintenance function of the NPC is evolutionarily conserved.

The importance of NPCs in genome stability is also underscored by studies showing that diverse DNA lesions relocate to NPCs. It has been reported that expanded CAG repeats cause replication fork collapse and relocation to the NPC, where the replication fork presumably restarts (Su et al., 2015). Other problematic DNA structures, like persistent DSBs (Nagai et al., 2008), eroded telomeres (Khadaroo et al., 2009) and R-loops (Penzo et al., 2023), also relocate to the NPC, implying a special repair-conducive environment ensuring genome integrity. Nup84, the C-terminal region of Nup1, and the NPC-associated SUMO-targeted ubiquitin ligase Slx5-Slx8 are required for relocation of diverse problematic DNA structures to the NPC (Nagai et al., 2008; Su et al., 2015; Churikov et al., 2016; Aguilera et al., 2020). The *nup84Δ*, *slx5Δ*, and *slx8Δ* mutants were reported to increase fragility, as well as contraction and expansion rates of CAG repeats (Su et al., 2015), suggesting that the NPC is either preventing breakage or facilitating repair of DNA damage at CAG repeats in a sumoylation-dependent manner. Nup84 is important for repairing DSBs via sister chromatid recombination (Gaillard et al., 2019), and it has been proposed that Nup84 helps suppress CAG expansions by regulating homologous recombination (Su et al., 2015). However, cells expressing Nup1 fused to LexA at its C-terminus, which is defective in the relocation of CAG repeats to the NPC, is not sensitive to DNA damaging agents and does not increase CAG repeat fragility or expansion, but does show a fourfold increase in CAG contraction rate (Aguilera et al., 2020). Thus, it is still unclear what role the NPC plays, and whether the observations with CAG repeats can be generalized to other STRs.

This study aims to clarify the importance of the relocation of STRs to the NPC and how it affects genome maintenance. We find that multiple expanded STRs relocate to the NPC and that the Nup1 C-terminus and Nup84 are both needed for this process. In *nup1ΔCt* cells, which express C-terminally truncated Nup1, relocation to the NPC is abrogated and contraction of STRs is increased, while repeat expansions are unaffected. Surprisingly, deletion of *NUP60*—which causes mislocalization of basket proteins Nup2, Mlp1, Mlp2, and Pml39 (Feuerbach et al., 2002; Niepel et al., 2013; Cibulka et al., 2022; Palancade et al., 2005), as well as the SUMO protease Ulp1 (Zhao et al., 2004)—does not affect relocation, yet *nup60Δ* cells have a greater STR contraction rate and are generally more genomically unstable than *nup1ΔCt* cells. Our results suggest that relocation to the NPC is likely a general mechanism to prevent contraction of disease-related STRs and is separable from the role of Nup60 in maintaining genomic stability.

## Results

### Short tandem repeat instability is sequence, length, and orientation dependent

STR instability positively correlates with the length of the repetitive sequence; the longer they are, the more unstable they get (Khristich and Mirkin, 2020). We first assessed the relationship between STR length and instability for different STRs, which are known fragile sites that are prone to breakage (Brown and Freudenreich, 2021). In case of a DNA break, cells mostly undergo error-free repair, but at a low frequency, they can engage in a mutagenic repair pathway leading to a gross chromosomal rearrangement (GCR). The rate of this rare event can be measured with a GCR assay, where the loss of metabolic markers placed on the left arm of chromosome V results in selectable growth (Putnam and Kolodner, 2010) (Fig. 1A). We inserted GAA/TTC, CAG/CTG, TGAAA/TTTCA repeats of varying length (up to 800 bp) next to the metabolic markers and measured the GCR rate. In this article, we write “CAG repeats” when referring to the CAG repeats being on the strand used as the template for lagging strand synthesis. We find that GCR rate exponentially increases with STR length (Fig. 1, B and C; Table S1). The exact relationship between GCR rate and length also depends on the sequence, which determines the type of noncanonical secondary structure formed, as well as its orientation (Khristich and Mirkin, 2020).

**Figure 1.**
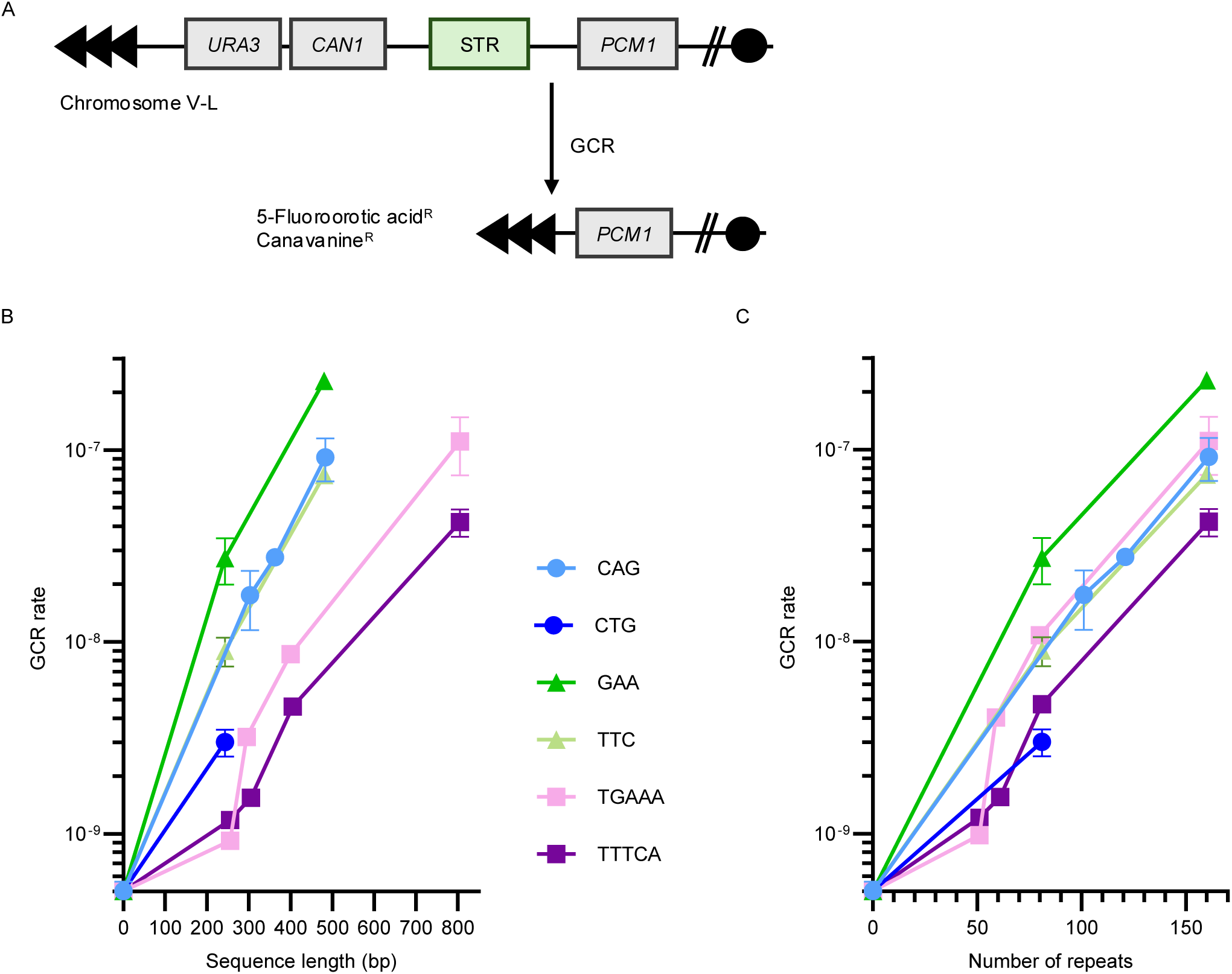
Short tandem repeat instability is dependent on the repetitive sequence, length and orientation. A) Schematic representation of the GCR assay. A GCR event that results in loss of the *URA3* and *CAN1* genes allows growth on the drugs 5-fluoroorotic acid and canavanine. *PCM1* is the most telomere-proximal essential gene on the left arm of chromosome V. B) GCR rate of different STRs plotted as a function of sequence length. C) GCR rate of different STRs plotted as a function of repeat number.

### Expanded CAG, CTG, TTTCA, TGAAA, GAA and TTC repeats relocate to the NPC in S/G2 phase

We next asked if relocation to the NPC is specific for CAG repeats (or hairpin-forming STRs) or if relocation is also observed with other STRs. Determining the length/instability relationship (Fig. 1; Table S1) allowed us to hypothesize which repeat lengths might be sufficient to see a relocation to the NPC. It was previously shown that 70 CAG repeats were sufficient to detect relocation to the NPC (Su et al., 2015). Thus, we decided to proceed with lengths that yield a GCR rate similar to or greater than that of (CAG)_70_. To track the location of the STR inside the nucleus, we used yeast strains harbouring a *lacO*/LacI-GFP system, in which a *lacO* array is bound by LacI-GFP 6.4 kb away from the repetitive sequence we are testing (Fig. 2A), as previously described (Su et al., 2015). To define the border of the nucleus, the nuclear envelope transmembrane protein Hmg1 was endogenously tagged with mCherry. We then scored the presence of the STR at the nuclear periphery (zone 1; Fig. 2B). As expected, with this zoning assay, we find that the (CAG)_73_ repeat sequence relocates to the nuclear periphery in S/G2 phase and not in G1 phase (Fig. 2D and S1; Table S2). In addition, (CTG)_67_, (TTTCA)_81_, (TGAAA)_81_, (GAA)_99_ and (TTC)_160_ do so as well.

**Figure 2.**
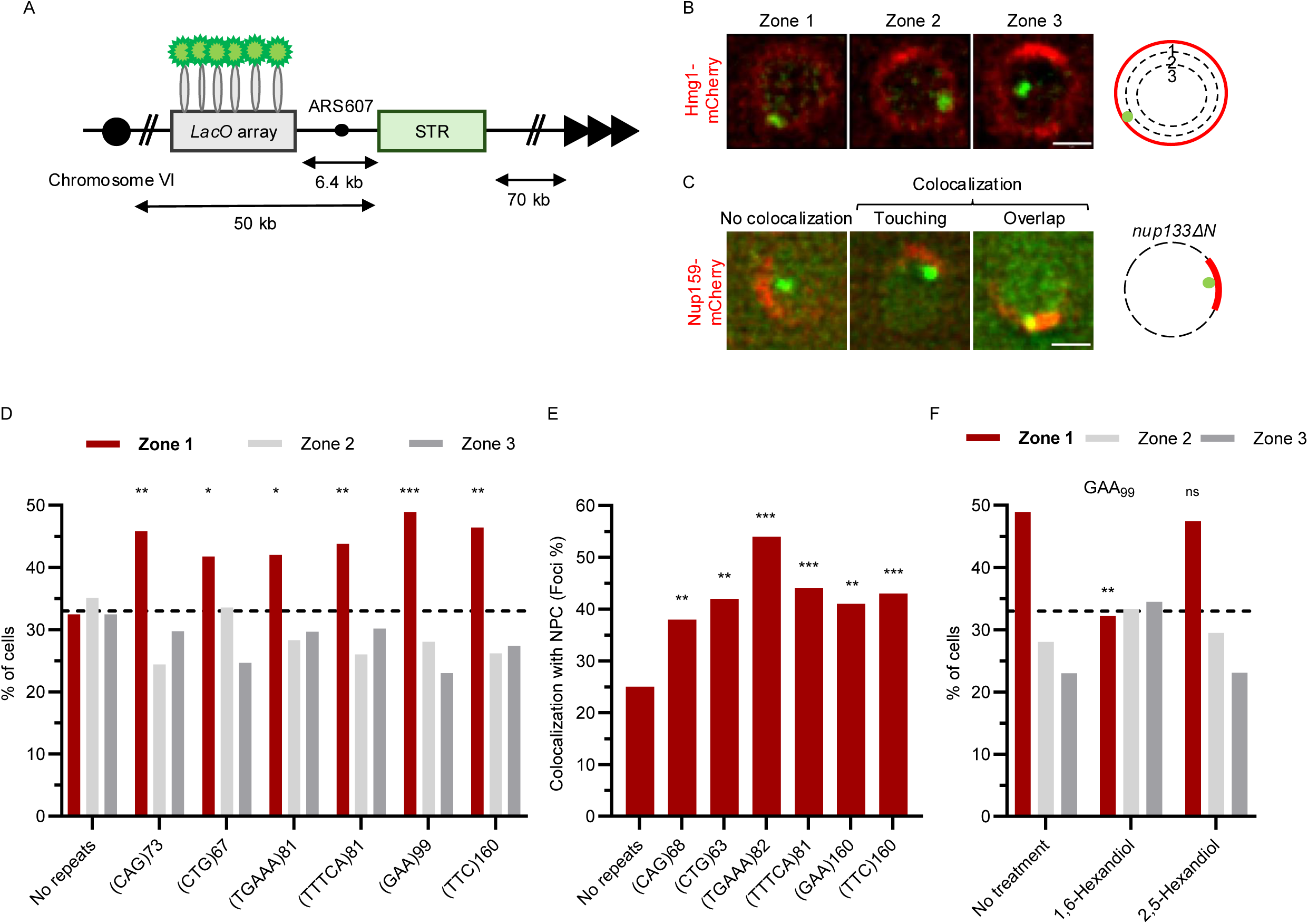
STRs relocate to the NPC in S/G2 phase. A) Organization of yeast chromosome VI used to score the localization of the STR inside the nucleus. The *lacO* array is bound by LacI-GFP 6.4 kb away from the STR; the nuclear envelope is marked by Hmg1-mCherry. B) Schematic and representative images of the localization of STRs inside the nucleus divided into three zones of equal volume. STR: green, LacI-GFP. Nuclear envelope: red, Hmg1-mCherry. Scale bar: 1 µm. C) Schematic and representative images assessing the localization of STRs relative to NPC clusters in *nup133ΔN*. When STRs and NPCs “touch” or “overlap”, they are categorized as co-localizing. Scale bar: 1 µm. D) STR localization in S/G2 phase cells, defined by the presence of a bud and a round nucleus, using the zoning assay depicted in panel B. (*) p<0.05, (**) p<0.01, (***) p<0.001 compared to no repeats, calculated using Fisher’s exact test. See Table S2 for raw numerical data and p-values. E) Co-localization of indicated STRs with NPC clusters measured and described as described in C. F) STR localization in S/G2 phase cells with (GAA)_99_ repeats, using the zoning assay depicted in panel B. (ns) non-significant, (**) p<0.01 compared to no hexanediol treatment, calculated using Fisher’s exact test. See Table S4 for raw numerical data and p-values.

The zoning assay does not differentiate between enrichment at the nuclear periphery in general or specifically at NPCs. To ensure the relocation is to NPCs, we made use of a mutant (*nup133ΔN*) in which NPCs are no longer evenly spaced along the nuclear envelope but clustered together (Fig. 2C). Importantly, this truncation of Nup133 does not cause defects in mRNA transport or DNA repair (Doye et al., 1994; Loeillet et al., 2005). In this mutant, we observe that the fluorescently labelled *lacO* array colocalizes significantly more with the clustered NPCs (visualized by the presence of Nup159-mCherry) when STRs are present (Fig. 2E; Table S3), confirming association with NPCs.

We noticed that the anchoring to NPCs might depend on weak hydrophobic interactions, as the STRs lose their enrichment at the nuclear periphery when exposed to 5% 1,6-hexanediol for 10 minutes, while this is not observed upon exposure to 2,5-hexanediol (Fig. 2F; Table S4). FG-Nups engage in weak hydrophobic interactions between their FG repeats and with nuclear transport receptors (NTRs), and previous work showed that a 10-minute exposure to 5% 1,6-hexanediol strongly affects the permeability barrier of NPCs and displaces several NTRs, while 2,5-hexanediol does less so (Riquelme Barrientos et al., 2023; Shulga and Goldfarb, 2003).

We conclude that all STRs tested localize to the NPC specifically, not only to the nuclear periphery, and that the association might rely on multivalent weak hydrophobic interactions, which could be provided by the NPC basket components.

### Relocation to the NPC is dependent on Nup84 and Nup1, but not Nup60

Next, we aimed to identify how the NPC might be involved in the relocation of the repeats. Previously, it was shown that relocation of CAG repeats to the NPC is abrogated in a mutant lacking Nup84 (*nup84Δ*) and upon truncation of the C-terminal 36 amino acids of Nup1 (*nup1ΔCt*) (Su et al., 2015; Aguilera et al., 2020). Since *nup84Δ* has been reported to cause mislocalization of the basket proteins Nup60 (partial delocalization into the nucleoplasm; Niño et al., 2016) and Mlp1/2 (nuclear foci; Niepel et al., 2013), we hypothesized that the basket is a key component for relocation of STRs to the NPC.

We assessed the effects of deleting/truncating subunits of the nuclear basket (Nup1 and Nup60) and Nup84 on NPC structure by examining the localization of endogenously GFP-tagged Nup1, Nup60, Nup84, Nup100 and Nup2 (Fig. 3A and S2) in *nup84Δ*, *nup60Δ* and *nup1ΔCt* mutants. Nup1 and Nup60 localization patterns are comparable in wild-type and *nup84Δ* strains, suggesting their correct incorporation into NPCs (Fig. 3A). As previously shown, the deletion of *NUP60* causes mislocalization of Nup2 (Fig. S5; Cibulka et al., 2022). Interestingly, the localization of Nup1-GFP is unchanged compared to wild-type cells (Fig. 3A), suggesting that, despite the absence of Nup2, Mlp1, Mlp2, and Pml39 (Feuerbach et al., 2002; Niepel et al., 2013; Cibulka et al., 2022; Palancade et al., 2005), Nup1 remains present at NPCs in *nup60Δ* cells. The truncation of the Nup1 C-terminus does not affect the other NPC proteins examined (Fig. 3A: Nup60, Nup84; Fig. S5: Nup2, Nup100), and it was previously shown that the localization of Nup1 itself is unchanged by this truncation (Pyhtila and Rexach, 2003). Altogether we conclude that the basket structure is significantly compromised when deleting *NUP84* and *NUP60*, but not when deleting 36 amino acids from the C-terminus of *NUP1* (Fig. 3B).

**Figure 3.**
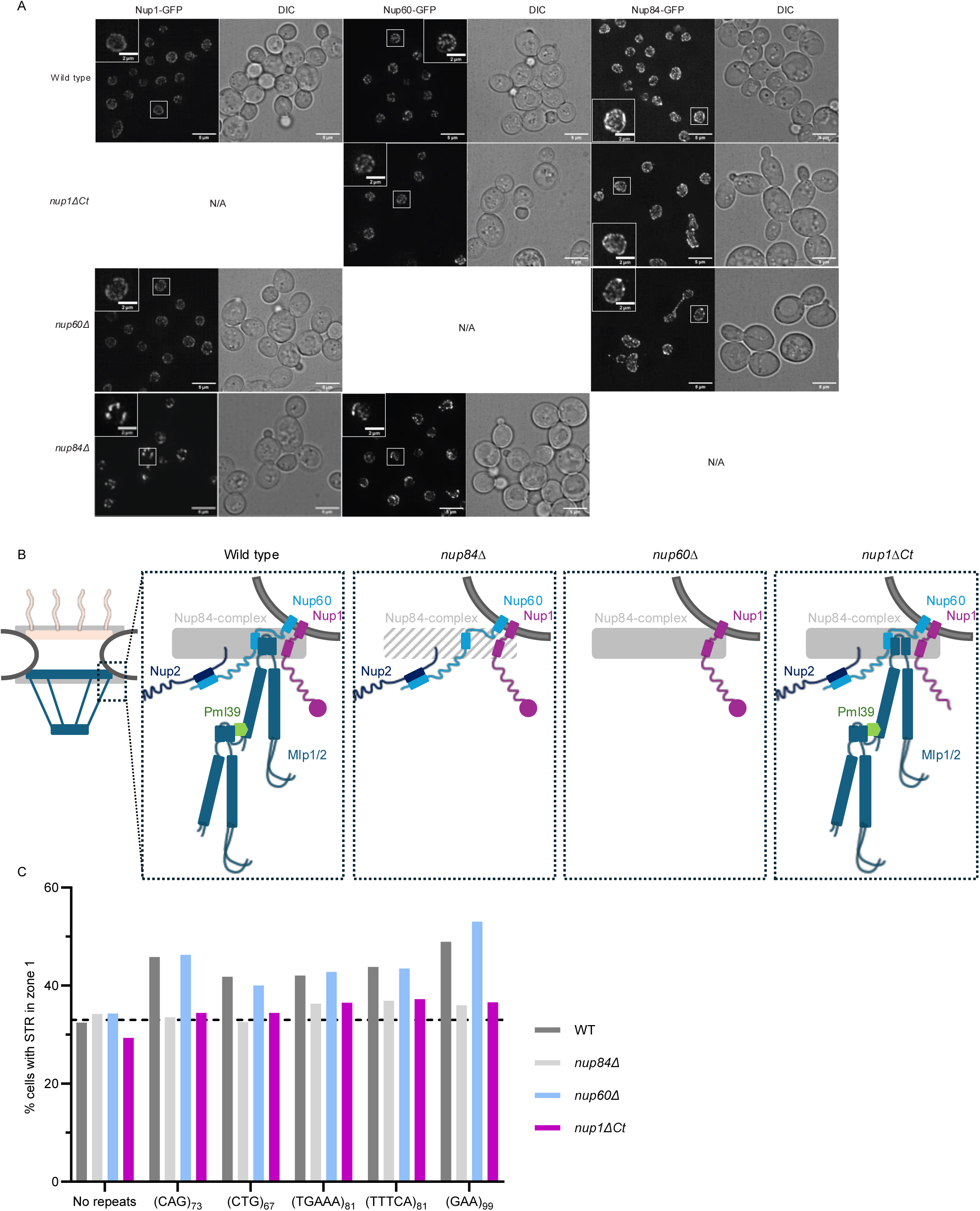
The relocation of STRs to the NPC is Nup84 and Nup1 dependent but Nup60 independent. A) Representative fluorescence microscopy images of the location of endogenously expressed Nup1-GFP, Nup60-GFP and Nup84-GFP in strains with the indicated genotypes. Images represent maximum intensity projections; scale bar: 5 μm (inset, 2 μm) B) Schematic representation of the NPC (left) and composition of the nuclear basket in wild type and indicated mutants as derived from A and (Dultz and Doye, 2025). Dark grey indicates the nuclear envelope; solid and striped light grey rectangles represents the outer ring in wild type and *nup84Δ*, respectively; circle at the end of Nup1 represents the last 36 amino acids at the C-terminal; wavy lines represent disordered sequences; green hexagon represents Pml39. C) Percentage of cells in S/G2 phase with GFP signal in zone 1. See Table S5 for raw numerical data and p-values. WT data are same as in Figure 2D.

We then tested the subnuclear localization of STRs in *nup84Δ*, *nup60Δ* and *nup1ΔCt* strains. Expanded CAG, CTG, TGAAA, TTTCA, and GAA repeats consistently exhibit greater enrichment at the nuclear periphery in S/G2-phase wild-type and *nup60Δ* cells than *nup84Δ* and *nup1ΔCt* cells (Fig. 3C and S3; Table S5). Thus, relocation to the NPC does not require Nup60, nor Nup2, Mlp1, Mlp2, Pml39, and Ulp1, as these proteins are not assembled at the NPC in the absence of Nup60 (Feuerbach et al., 2002; Zhao et al., 2004; Palancade et al., 2005; Palancade et al., 2007; Niepel et al., 2013; Cibulka et al., 2022). Furthermore, the observation that Nup1 and Nup84 are not mislocalized in *nup84Δ* and *nup1ΔCt* cells, respectively (Fig. 3A), suggests that Nup84 and the C-terminus of Nup1 are acting independently to promote relocation to the NPC.

### Nup1 C-terminus is not required for direct repeat recombination

We next asked which DNA repair pathway might be impacted in the *nup60Δ, nup84Δ* and *nup1ΔCt* mutants. It has been proposed that the pore is a docking site for DNA lesions and a location where noncanonical repair pathways take place (Freudenreich and Su, 2016; Simon et al., 2024). It has also been suggested that the NPC prevents telomeric sister chromatid recombination (Aguilera et al., 2020) and that collapsed replication forks at STRs could be repaired via homologous recombination (Su et al., 2015). We therefore tested if the NPC, and especially the *nup1ΔCt* mutant, has an impact on homologous recombination using a direct repeat recombination assay that detects recombination between two *leu2* heteroalleles (Smith and Rothstein, 1999; Fig. 5A). Consistent with prior observations (Gaillard et al., 2019), the *nup84Δ* mutant exhibits a tenfold increase in recombination rate compared to wild type, but *nup1ΔCt* has no significantly different rate (Fig. 4C), suggesting that relocation to the NPC does not impact homologous recombination. We also modified the direct repeat recombination assay by inserting (GAA)_160_ or (TTC)_160_ repeats in between the *leu2* heteroalleles (Fig. 4B). We observed a ∼15-fold increase in the recombination rate in the *nup84Δ* mutant, slightly higher than the tenfold change without STRs, while the *nup1ΔCt* mutant has no statistically significant effect, indicating a negligible effect of the mutant on recombination (Fig. 4D).

**Figure 4.**
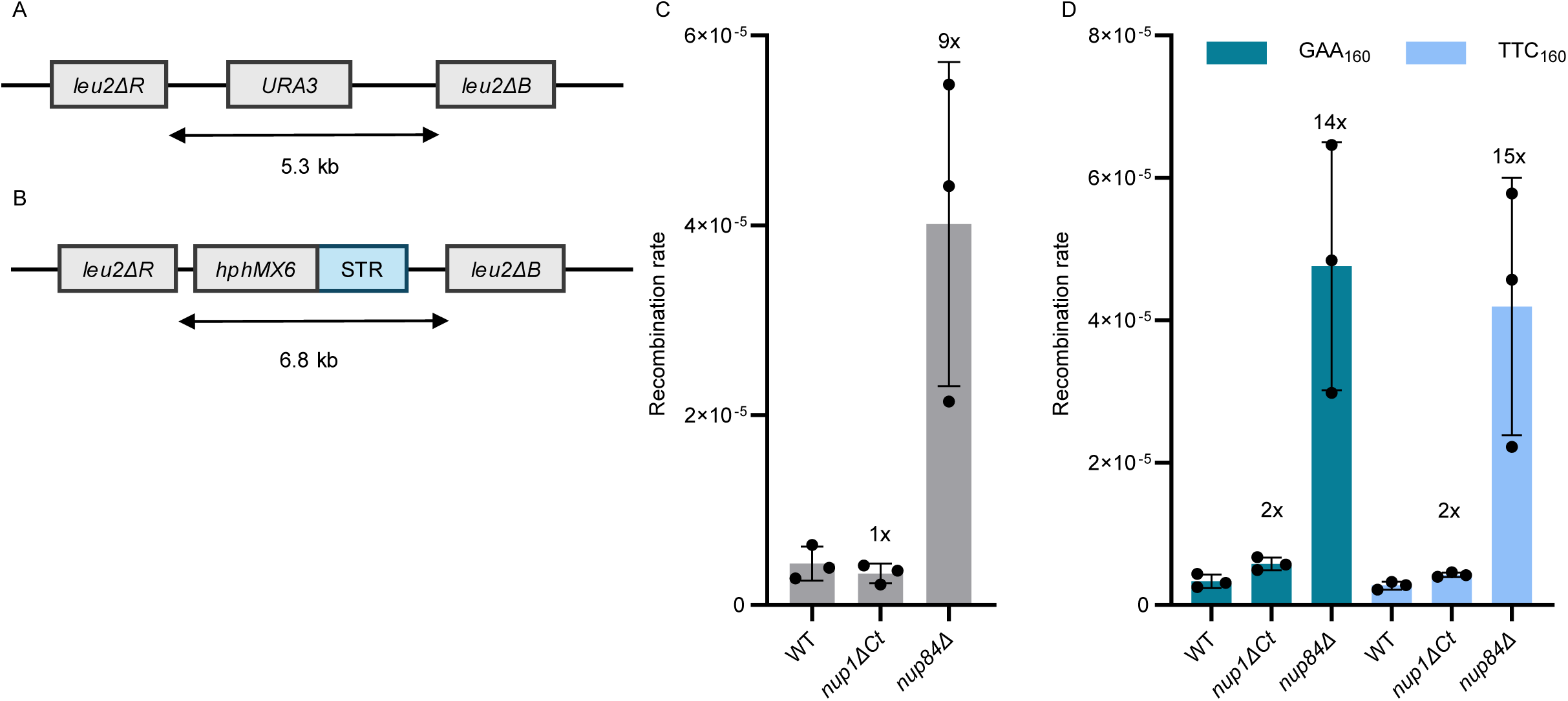
Homologous recombination are not affected in the *nup1ΔCt* mutant. A) Schematic representation of the *leu2* direct repeat recombination assay. B) Schematic representation of the *leu2* direct repeat recombination assay with STRs inserted between the *leu2* heteroalleles. C) Recombination rate, measured using the assay in A, of strains of the indicated genotypes. Numbers above bars indicate fold change compared to WT. D) Recombination rate, measured using the assay in B (with GAA_160_ or TTC_160_), of strains of the indicated genotypes.

**Figure 5.**
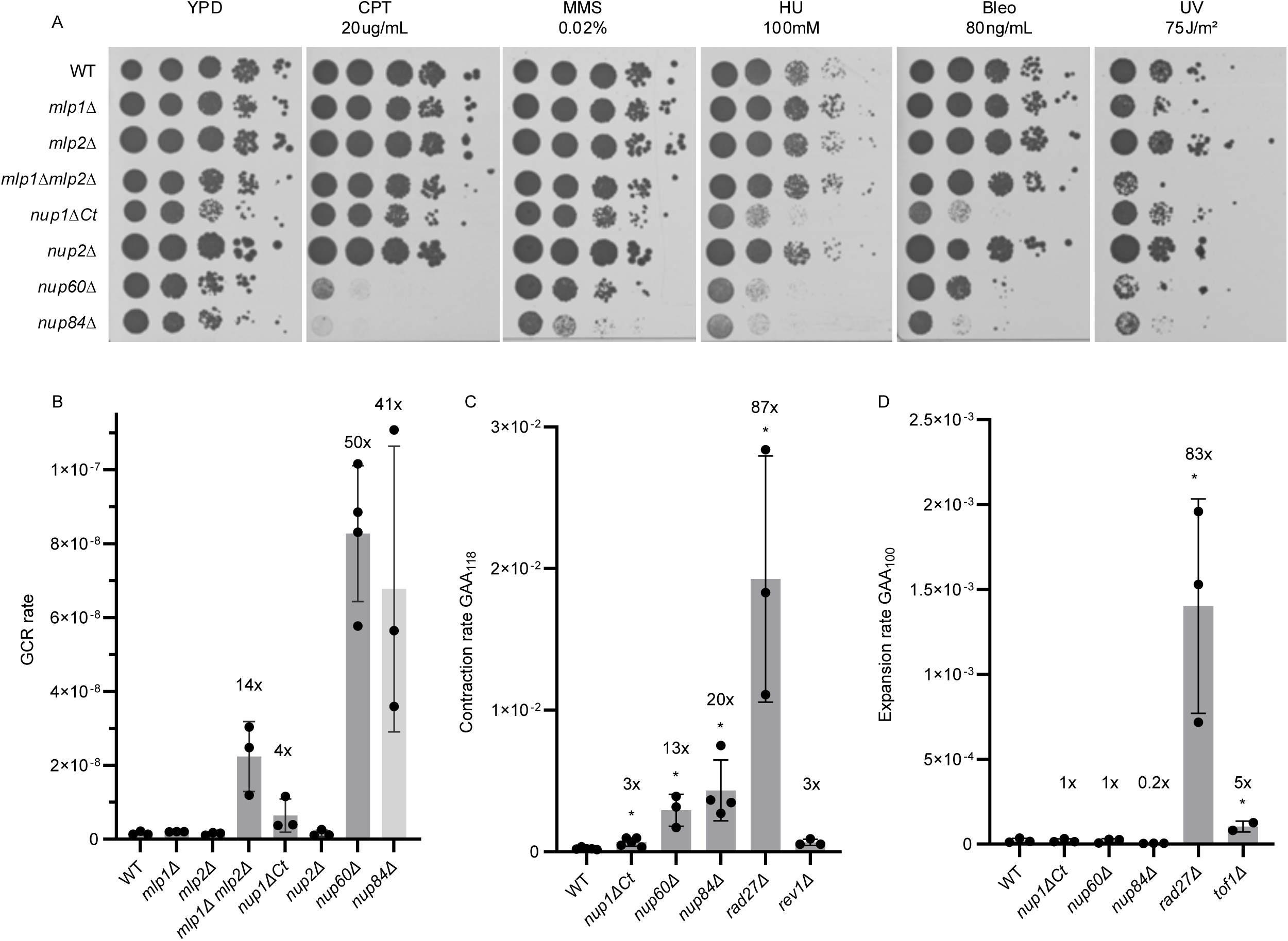
Genome stability is more strongly compromised in *nup84Δ* and *nup60Δ* than in *nup1ΔCt*. A) Tenfold serial dilutions of strains with the indicated genotypes were spotted on YPD plates with different DNA damaging agents. CPT: camptothecin, MMS: methyl methanesulfonate, HU: hydroxyurea, Bleo: bleomycin, UV: ultraviolet radiation. B) GCR rate of strains with the indicated genotypes. Numbers above bars indicate fold change compared to WT. C) GAA_118_ contraction rates for strains of the indicated genotypes. Numbers above bars indicate fold change compared to WT. (*) p <0.001 with Mann-Whitney U test. Rates and p-values can be found in Table S7 D) GAA_100_ expansion rates for strains of the indicated genotypes. (*) p <0.001 with Mann-Whitney U test. Rates and p-values can be found in Table S8.

### Genome stability is more strongly compromised in *nup84Δ* and *nup60Δ* than in *nup1ΔCt*

Nup84 and Nup60 are required for resistance to DNA damaging agents (Bennett et al., 2001; Chang et al., 2002; Hanway et al., 2002; Gaillard et al., 2019), and *nup84Δ* and *nup60Δ* strains have elevated levels of DNA repair foci (Palancade et al., 2007), GCR (Putnam et al., 2012), and spontaneous recombination (Alvaro et al., 2007; Palancade et al., 2007; Gaillard et al., 2019). *NUP84* and *NUP60* also have significant genetic interactions with genes important for genome stability (Loeillet et al., 2005; Palancade et al., 2007; Nagai et al., 2008). However, a role of Nup1 in genome stability has not been reported, although this may be due to *NUP1* being an essential gene that is absent from genome-wide studies using the yeast knockout collection (Davis and Fink, 1990; Giaever et al., 2002). To assess the importance of the basket versus Nup84 for genome maintenance, we again compared *nup1ΔCt*, *nup60Δ* and *nup84Δ*.

We find, as previously shown, that *nup84Δ* is sensitive to all DNA damaging agents tested (Fig. 5A) and shows a large increase in GCR rate (41-fold higher than wild type; Fig. 5B; Table S6). The *nup60Δ* mutant also exhibits a large increase in GCR rate (50-fold), and is sensitive to most of the DNA damaging agents tested, namely camptothecin (CPT), hydroxyurea (HU), bleomycin, and ultraviolet (UV) radiation. By contrast, despite its relocation defect, we find that *nup1ΔCt* is only sensitive to HU and bleomycin (Fig. 5A), and has a mild—as compared to other mutants with elevated rates of GCR (Putnam et al., 2012)—fourfold increase in GCR rate compared to wild type (Fig. 5B; Table S6). Our results are consistent with a previous study showing that a *nup1-LexA* fusion mutant, which is also defective in relocation to the NPC, is not sensitive to UV, MMS, or HU (Aguilera et al., 2020). The difference in HU sensitivity between *nup1ΔCt* and *nup1-LexA* may be due to the latter being a less defective allele.

To more directly address the stability of STRs in the *nup1ΔCt*, *nup60Δ* and *nup84Δ* strains, we performed contraction and expansion assays (Fig. 5, C and D; Tables S7 and S8). The contraction and expansion assays involve the insertion of an artificial intron into the *URA3* gene, and takes advantage of the fact that introns longer than ∼1.1 kb cannot be efficiently spliced in *S. cerevisiae* (Radchenko et al., 2018). In the contraction assay, GAA repeats were placed within the intron such that the total length of the intron is greater than this size threshold; therefore, the intron cannot be efficiently spliced unless a contraction of the GAA sequence occurs. Conversely, in the expansion assay, the intron was designed to be shorter than the threshold and can be spliced efficiently unless an expansion event occurs. We included a strain deleted for *RAD27* because it has been shown to have greatly elevated levels of both GAA repeat contraction and expansion (Khristich et al., 2020; Zhang et al., 2012). We also included *rev1Δ* and *tof1Δ*, which have been reported to have elevated levels of GAA contraction and expansion, respectively, but not as elevated as in *rad27Δ* (Khristich et al., 2020; Zhang et al., 2012). We find that *nup84Δ*, *nup60Δ*, and *nup1ΔCt* increase GAA repeat contraction rate 20-, 13-, and threefold, respectively; (Fig. 5C; Table S7), but none increase GAA expansion rate (Fig. 5D; Table S8). While more modest than that of *rad27Δ*, *nup84Δ*, and *nup60Δ*, the threefold effect of *nup1ΔCt* on GAA repeat contraction rate is similar to that of *rev1Δ* and *pol32Δ*; Rev1 and Pol32 have previously been proposed to act together to suppress GAA repeat contractions during lagging strand synthesis (Khristich et al., 2020). Our results are also consistent with a previous study showing that a *nup1-LexA* fusion causes a fourfold increase in CAG repeat contractions and no statistically significant effect on CAG repeat expansions (Aguilera et al., 2020).

Taken together, we conclude that the C-terminus of Nup1 is required for anchoring STRs to NPCs to prevent repeat contraction. In addition, genome stability is more strongly compromised in *nup60Δ* and *nup84Δ* cells than in *nup1ΔCt* cells. Since relocation to NPCs is abrogated in *nup1ΔCt* but not in *nup60Δ*, relocation to NPCs is separable from other genome stability roles of the NPC.

## Discussion

Expanded CAG repeats were previously reported to relocate to the NPC during S/G2 phase, requiring both Nup84 and the C-terminus of Nup1 (Su et al., 2015; Aguilera et al., 2020), but it was unclear whether other STRs involved in repeat expansion diseases would do so as well. Here, we show that expanded CTG, TTTCA, TGAAA, GAA and TTC repeats also relocate to the NPC in a Nup84- and Nup1-dependent manner, suggesting a general phenomenon. A full deletion of *NUP84* perturbs the structure and function of the NPC outer ring (Fernandez-Martinez et al., 2012), causes mislocalization of the basket components Mlp1 and Mlp2 (Niepel et al., 2013), and exhibits a broad genome instability phenotype, complicating its use to specifically examine relocation to the NPC. By contrast, the *nup1ΔCt* truncation does not affect the localization of NPC proteins (Nup100, Nup84, Nup60, Nup2) and has a much milder effect on genome stability. With respect to STRs, the *nup1ΔCt* mutant exhibits an increase in repeat contraction and no impact on repeat expansion. We also focused on Nup60, since its deletion results in mislocalization of all nuclear basket proteins (Feuerbach et al., 2002; Palancade et al., 2005; Niepel et al., 2013; Cibulka et al., 2022) except Nup1 (Fig. 3A). The *nup60Δ* deletion causes greater sensitivity to DNA damaging agents and larger increases in the rates of GCR and GAA repeat contraction than *nup1ΔCt*. However, we find that the *nup60Δ* deletion does not affect relocation of STRs to the NPC. It has been suggested that relocation of expanded STRs to the NPC is important to prevent repeat expansion, contraction, and fragility (Su et al., 2015; Whalen et al., 2020). However, *mrc1^AQ^* and *cep3^S575A^* mutants are relocation defective, yet repeat fragility is largely unaffected (Gellon et al., 2019; Maclay et al., 2025). Furthermore, our study and a prior publication (Aguilera et al., 2020) have found that the *nup1ΔCt* and *nup1-LexA* mutants are also relocation defective and display an increase in STR contraction, but not expansion. Taken together, these findings suggest that the role of the NPC in maintaining genome stability is only partially dependent on the ability to relocate damaged or difficult-to-replicate DNA to the NPC.

The phenotype of the *nup1ΔCt* mutant is striking, but presently not fully understood. Nup1, like other FG-Nups, encodes weak binding sites for nuclear transport receptors (NTRs), yet the C-terminal domain of Nup1 is special, as it interacts with the NTR Kap95 over 200-fold more strongly than typical FG-Nups (Pyhtila and Rexach, 2003). Crystallography showed that the amino acid regions 975–987 and 1004–1009 of Nup1 bind extensively with Kap95, explaining the higher affinity (Liu and Stewart, 2005). Even though these regions are retained in the Nup1ΔCt mutant, which only lacks amino acids 1040–1076, biochemical studies do show impaired Kap95 binding, and, in cells, Kap95-dependent nuclear import is reduced, and Kap95 and Kap60 are partially mislocalized (Pyhtila and Rexach, 2003). We observe that 1,6-hexandiol, which disrupts the permeability barrier and displaces NTRs from the NPC (Riquelme Barrientos et al., 2023), also prevents the localization of STRs to the NPC (Fig. 2F). This may support previous speculations that NTRs might be involved (Aguilera et al., 2020), but future research is needed.

The mild genomic instability phenotypes observed in *nup1ΔCt* and *nup1-LexA* can be explained by the possibility that when STRs fail to relocalize efficiently to the NPC, repair may still proceed at alternative intranuclear sites. While Nup1 is required for the relocation of STRs to the NPC, Nup60 could function downstream as a platform for processing expanded STRs. Indeed, the SUMO protease Ulp1 is normally anchored at the nuclear basket through Nup60 and Mlp1/2, and in *nup60Δ* strains, Ulp1 becomes mislocalized and reduced in abundance, leading to defects in SUMO homeostasis (Zhao et al., 2004; Palancade et al., 2007). Furthermore, the genetic interaction profile of *NUP60* is similar to those of *SLX5* and *SLX8*, which encode subunits of a heterodimeric SUMO-targeted ubiquitin ligase (STUbL) that associates with NPCs (Nagai et al., 2008), and proteasomes are tethered to the nuclear basket (Albert et al., 2017), suggesting that Nup60 may help mediate the removal of ubiquitinated proteins at stalled forks to enable replication restart. Together, these observations could explain why *nup60Δ* exhibits stronger defects in genome stability than the *nup1ΔCt* mutant; relocating expanded STRs to the NPC without downstream Ulp1- and STUbL-dependent processing may be more detrimental than failing to relocate them at all.

Despite their mild overall phenotypes, *nup1ΔCt* (this study) and *nup1-LexA* (Aguilera et al., 2020) mutants nonetheless display a measurable and consistent increase in repeat contraction. We propose that NPC anchoring may help suppress contractions by stabilizing stalled replication intermediates or preventing aberrant processing pathways. This raises the possibility that STRs may have evolved mechanisms to maintain their length, analogous to telomeric repeats, and that NPC association contributes to this stability. Our data suggest that this contraction phenotype does not depend on homologous recombination. One possibility is that break-induced replication (BIR) is involved. BIR has been implicated in large-scale expansions of CAG repeats (Kim et al., 2017) as well as transcription-induced expansions of GAA repeats (Neil et al., 2018). Eroded telomeres also relocate to NPCs, where they can be extended by BIR (Khadaroo et al., 2009; Aguilera et al., 2022). Furthermore, strains lacking *NUP84* and *NUP60* are defective in BIR (Chung et al., 2015; Horigome et al., 2016; Oshidari et al., 2018; Liu et al., 2025). Future work will be required to determine if and how BIR at NPCs prevent STR contraction.

In conclusion, our findings provide novel insight into the role of the NPC in maintaining genome stability. Since both genome instability and NPC dysfunction are linked to aging, cancer, and neurological disorders, elucidating the role of the NPC could open up new perspectives on the molecular origins of these conditions.

## Materials and methods

### Cloning of repetitive sequences

Cloning and expansion of repetitive sequences was performed in vector pGC542 as described (Williams and Coster, 2024). Briefly, the synthetic DNA fragment PacI-BsaI-STR_n_-BsmBI-NotI was cloned in pGC542 between PacI and NotI restriction sites. The resulting plasmid was then digested with BsmBI/NotI to obtain the vector, containing STR_n_, and with BsaI/NotI to obtain the insert containing STR_n_. Subsequent ligation of the vector and the insert yielded a plasmid containing twice as many repeats (PacI-BsaI-STR_2n_-BsmBI-NotI). The cycle of digestion and ligation was repeated to expand the repeats until a tract of uninterrupted repeats of the desired length was obtained. The STR sequences were then subcloned via PacI/NotI digestion and ligation in vectors pDN7.8 or pDN12.1 for integration at the *PRB1* locus (GCR assay), in vectors pSO31 or pSO32 for integration at the *HIS2* locus (zoning assay), and in vector pSO81 or pSO82 for integration at the *ARS306* locus (expansion and contraction assay, respectively).

Verification of the repeat size was done when possible by Sanger sequencing (Gietz and Schiestl, 2007), and otherwise by digestion of flanking restriction sites PacI/NotI followed by TBE-PAGE on an 8% acrylamide gel and staining with SYBR Safe (Invitrogen) (Casas-Delucchi et al., 2022). The cloned plasmids were amplified in NEB® Stable Competent *E. coli* grown at 30°C to slow down replication and maintain repeat size length.

Plasmid pSO53 for CRISPR/Cas9 genome editing was constructed with the MoClo-YTK (Lee et al., 2015) as described (Novarina et al., 2022). NUP133-sgRNA (TTGGGAAAGTTAGTATGCACCT) was cloned in pYTK050 via BsmBI Golden Gate assembly of partially overlapping oligonucleotides. The resulting “sgRNA part plasmid” was assembled with plasmids pYTK003, pYTK068 and pYTK095 via BsaI Golden Gate assembly to obtain a “sgRNA cassette plasmid”. Finally, the sgRNA cassette was combined with Cas9 in a multigene vector via BsmBI Golden Gate assembly of plasmids pYTK-DN1 + sgRNA cassette plasmid + pYTK-DN2 + pYTK-DN5.

### Yeast strains

Yeast strains used for this study are listed in Table S10. All yeast transformations were checked for correct insertion by PCR and for repeat size by Sanger sequencing (Gietz and Schiestl, 2007). Gene deletions were performed by PCR-mediated gene replacement via marker amplification from the yeast knockout collection (Giaever et al., 2002). *nup1ΔCt* was created by inserting a stop codon and a kanMX cassette to remove the 36 C-terminal amino acids.

The *nup133ΔN* mutation (*nup133^Δ2-300^*) was made using CRISPR/Cas9 technology (Novarina et al., 2022) by co-transforming yeast with the sgRNA- and Cas9-containing plasmid pSO53 and the repair fragment. The repair fragment was obtained by PCR amplification of partially overlapping oligonucleotides gRNA nup133N fwd (GACTTTGGGAAAGTTAGTATGCACCT) and gRNA nup133N rev (AAACAGGTGCATACTAACTTTCCCAA). After verification of genome editing via PCR, the plasmid was removed by growing the strain in non-selective media.

### Zoning assay

Yeast cells were grown overnight in 4 ml SD complete, diluted in the morning (0.1 ml in 4 ml SD complete), and again in the evening (0.005 ml in 10 ml SD complete). The following morning, the optical density was measured and cells were microscoped when in log phase (OD_600_ between 0.4 and 0.8). A single DIC image along with 18 GFP and 18 mCherry images obtained at 0.2-µm intervals along the z-axis were captured per frame. The distance from the GFP locus to the nearest mCherry signal was measured as well as the diameter of the nucleus, as previously described (Meister et al., 2010). At least 100 cells per strain were analysed. Cell cycle phase was determined by bud morphology: unbudded cells were scored as being in G1 phase whereas budded cells with a round nucleus were scored as being in S/G2 phase. For the hexanediol experiment, cells were incubated in 5% 1,6-hexanediol or 2,5-hexanediol for 10 min at 30°C, followed by an additional 5 min incubation on a ConA-coated coverslip prior to imaging. Imaging was peformed using a Deltavision Elite (Applied Precision) microscope equipped with an Olympus UPlanSApo 100x (NA1.4) oil immersion objective using InsightSSIT Solid State Illumination. Detection was performed by an EDGE sCMOS5.5 camera. Images were deconvolved using softWoRx before ImageJ analysis.

### Gross chromosomal rearrangement assay

Fluctuation tests for the quantification of GCR rates was performed essentially as described (Putnam and Kolodner, 2010; Yao et al., 2024) using the method of the median (Lea and Coulson, 1949). Each experiment was performed at least three times.

### Expansion and contraction assay

Expansion and contraction assays were performed as previously described (Radchenko et al., 2018). Strains were streaked on YPD supplemented with uracil for 40 h (64 h for *nup84Δ* (SOY213), due to its growth defect). Single colonies were picked and added to 200 µl water, followed by five 1:10 serial dilutions. The 10^-5^ dilution was used to plate on YPD plates, and the 10^-1^ to 10^-3^ dilutions for the selective plates. The selective plates contained 2 g/l of 5-fluoroorotic acid for the expansion assay, while for the contraction assay, SD-ura plates were used. After 4 d in the 30°C incubator, the colonies were counted and at least 4 colonies per plate were size analyzed by PCR. For the contraction assay, the PCR step was left out since 100% of colonies growing on selection plates are due to contraction. Expansion or contraction rate was calculated with the method of the median (Lea and Coulson, 1949). Each experiment was performed at least three times.

### Homologous recombination assay

The *leu2* direct repeat recombination assays were performed essentially as previously described (Novarina et al., 2020). The strains were streaked for single colonies on a YPD plate and incubated at 30°C for 2 d (3 d for *nup84Δ*). Single colonies were inoculated in 4 ml YPD and grown to saturation for 24 h at 30°C. Dilutions were plated on YPD (10^5^) and SD-leu (10^-1^). Cells were counted after 3 d of incubation at 30°C. Recombination rate was calculated with the method of the median (Lea and Coulson, 1949). Each experiment was performed three times.

## Acknowledgments

We thank Marie-Noëlle Simon and Sergei Mirkin for providing yeast strains and plasmids. This work was supported by a Vici grant (VI.C.192.031 to LMV) and an ENW M-2 grant (OCENW.M20.068 to LMV and MC) from the Dutch Research Council.

## Supplemental material

**Figure S1.**
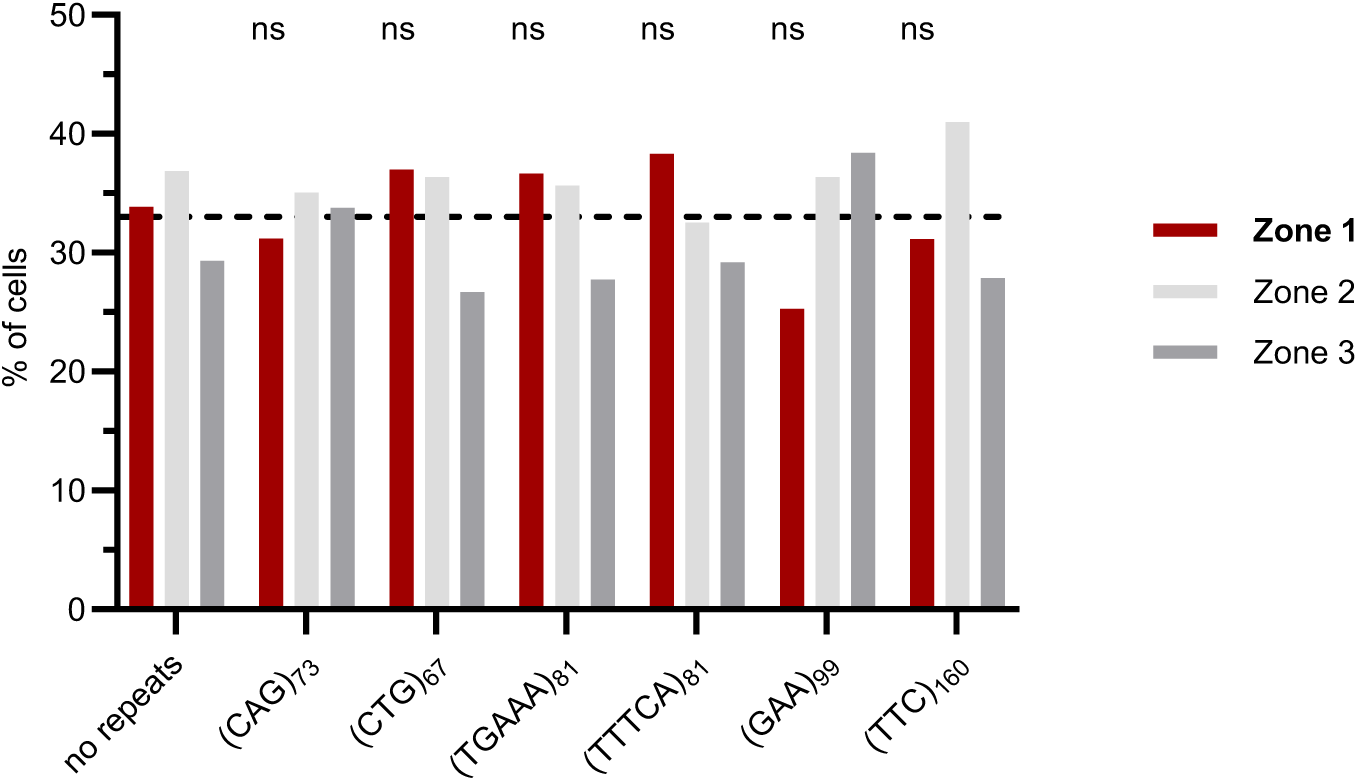
Subnuclear localization of the indicated STRs in G1 phase cells, defined by the absence of a bud, as determined by the zoning assay (complement to Fig. 2D). ns: not significant compared to cells without STRs inserted (zone 1) by Fisher’s exact test.

**Figure S2.**
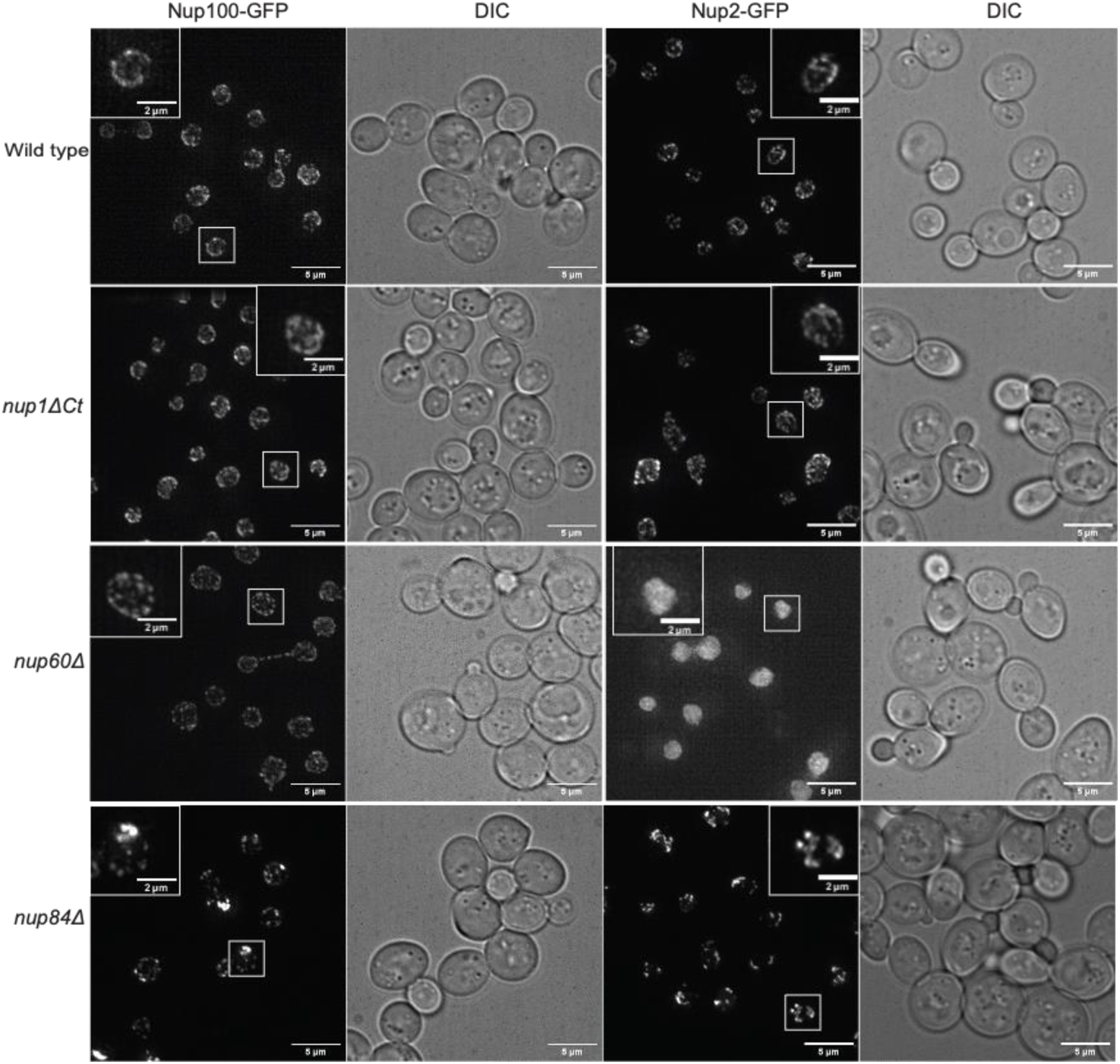
Representative fluorescence microscopy images of endogenously expressed Nup100-GFP and Nup2-GFP in strains of the indicated genotypes. Images represent maximum intensity projections; scale bar: 5 μm (inset, 2 μm)

**Figure S3.**
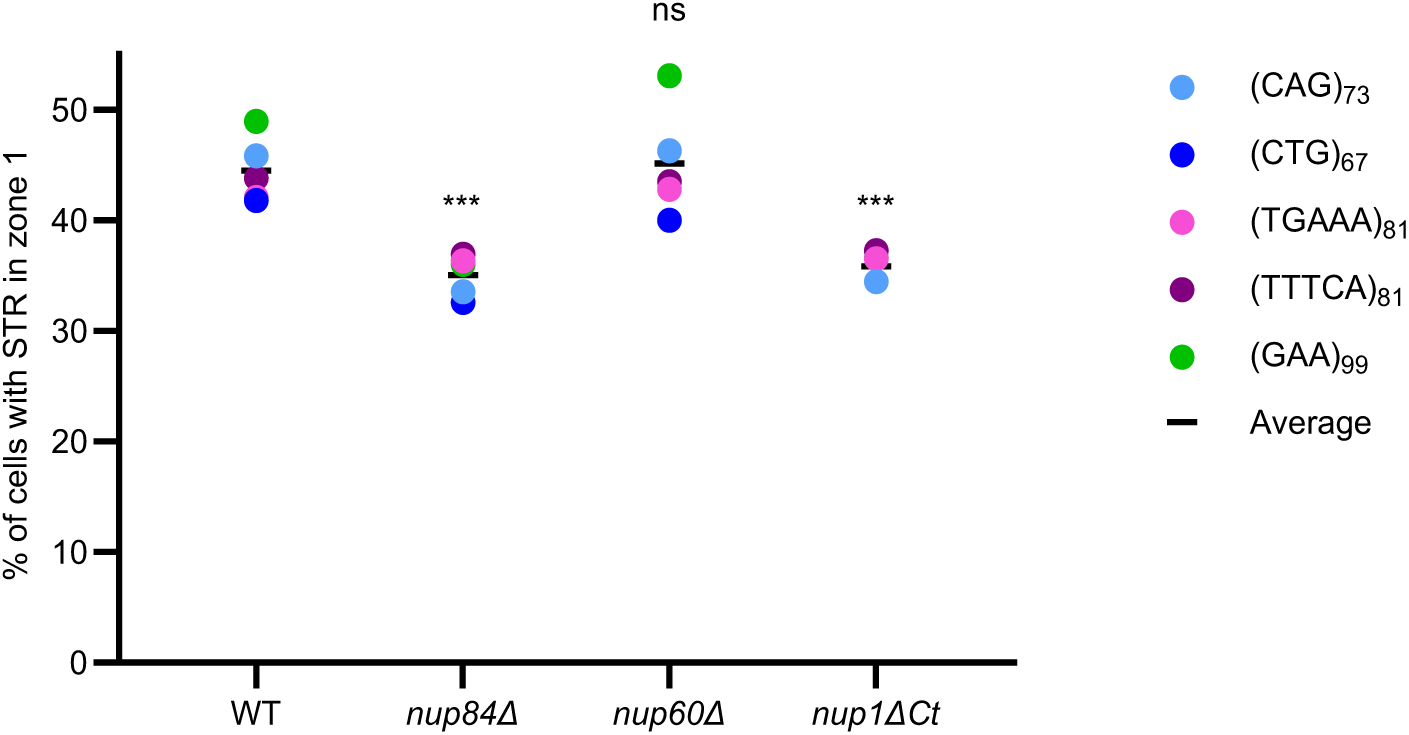
Percentage of cells in S/G2 phase with GFP signal in zone 1. Different representation of data from Fig. 3C, with average percentage of all STRs indicated. p-values were calculating using a pooled mixed-effects logistic regression with dataset included as a random effect. *nup84Δ*: odds ratio (OR) = 0.674, 95% confidence interval (CI) [0.551, 0.823], p = 0.0001; *nup60Δ*: OR = 1.018, 95% CI [0.842, 1.230], p = 0.857, *nup1ΔCt*: OR = 0.696, 95% CI [0.575, 0.842], p = 0.0002.

**Table S1.**
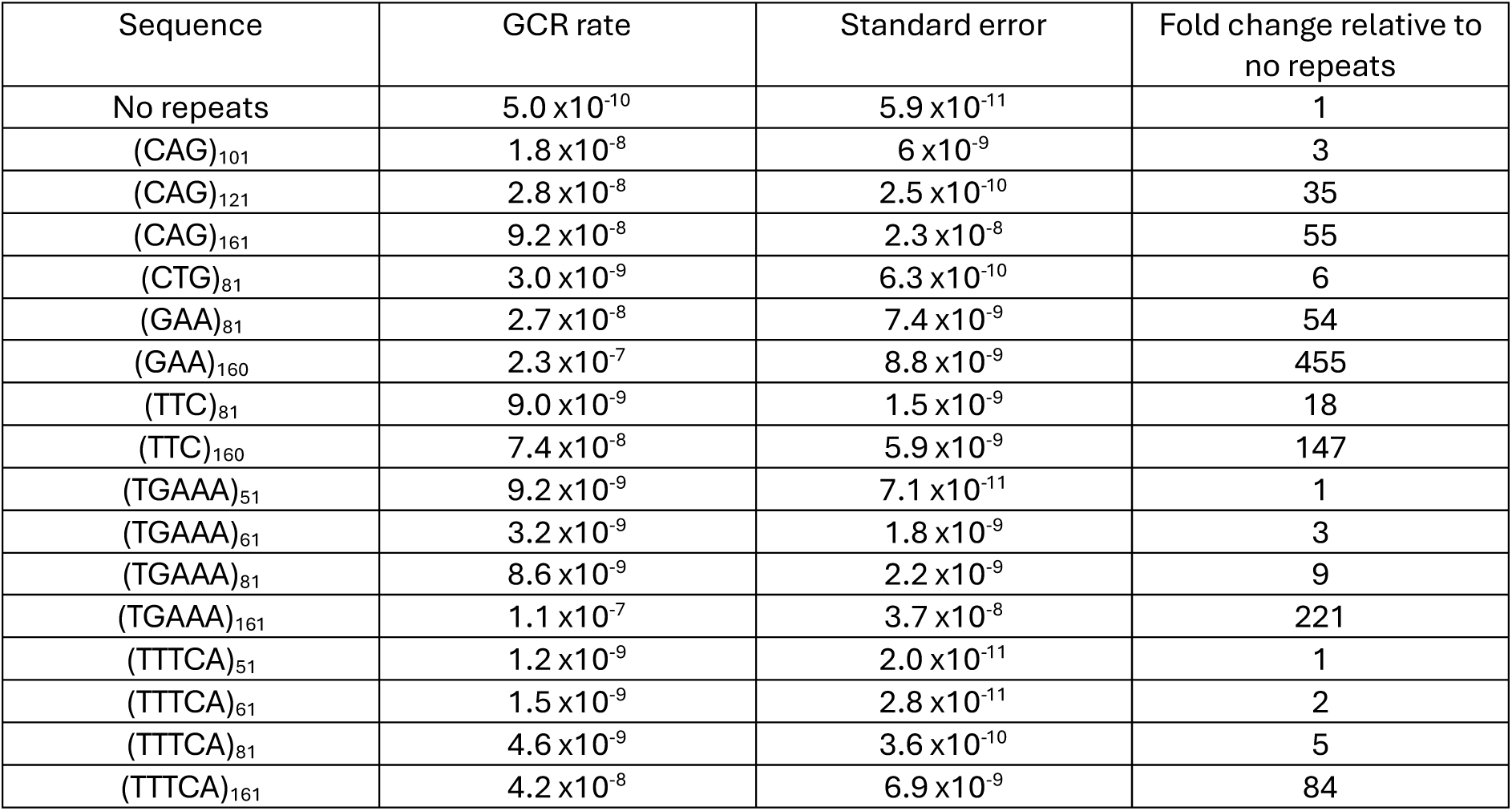
GCR rates for Figure 1B and C.

| Sequence | GCR rate | Standard error | Fold change relative to no repeats |
| --- | --- | --- | --- |
| No repeats | $5.0 \times 10^{-10}$ | $5.9 \times 10^{-11}$ | 1 |
| (CAG)<sub>101</sub> | $1.8 \times 10^{-8}$ | $6 \times 10^{-9}$ | 3 |
| (CAG)<sub>121</sub> | $2.8 \times 10^{-8}$ | $2.5 \times 10^{-10}$ | 35 |
| (CAG)<sub>161</sub> | $9.2 \times 10^{-8}$ | $2.3 \times 10^{-8}$ | 55 |
| (CTG)<sub>81</sub> | $3.0 \times 10^{-9}$ | $6.3 \times 10^{-10}$ | 6 |
| (GAA)<sub>81</sub> | $2.7 \times 10^{-8}$ | $7.4 \times 10^{-9}$ | 54 |
| (GAA)<sub>160</sub> | $2.3 \times 10^{-7}$ | $8.8 \times 10^{-9}$ | 455 |
| (TTC)<sub>81</sub> | $9.0 \times 10^{-9}$ | $1.5 \times 10^{-9}$ | 18 |
| (TTC)<sub>160</sub> | $7.4 \times 10^{-8}$ | $5.9 \times 10^{-9}$ | 147 |
| (TGAAA)<sub>51</sub> | $9.2 \times 10^{-9}$ | $7.1 \times 10^{-11}$ | 1 |
| (TGAAA)<sub>61</sub> | $3.2 \times 10^{-9}$ | $1.8 \times 10^{-9}$ | 3 |
| (TGAAA)<sub>81</sub> | $8.6 \times 10^{-9}$ | $2.2 \times 10^{-9}$ | 9 |
| (TGAAA)<sub>161</sub> | $1.1 \times 10^{-7}$ | $3.7 \times 10^{-8}$ | 221 |
| (TTTCA)<sub>51</sub> | $1.2 \times 10^{-9}$ | $2.0 \times 10^{-11}$ | 1 |
| (TTTCA)<sub>61</sub> | $1.5 \times 10^{-9}$ | $2.8 \times 10^{-11}$ | 2 |
| (TTTCA)<sub>81</sub> | $4.6 \times 10^{-9}$ | $3.6 \times 10^{-10}$ | 5 |
| (TTTCA)<sub>161</sub> | $4.2 \times 10^{-8}$ | $6.9 \times 10^{-9}$ | 84 |

**Table S2.**
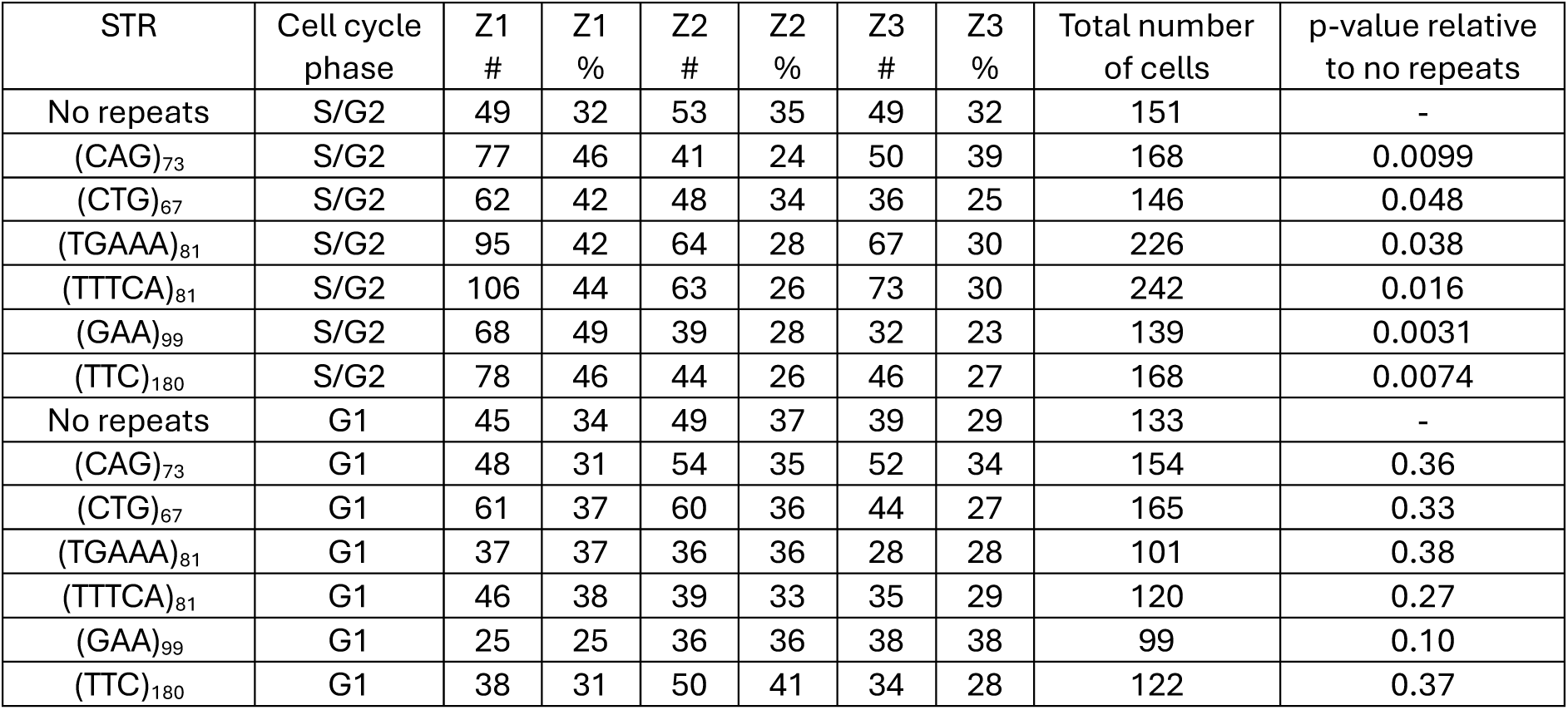
Zoning analysis for Figures 2D and S1.

**Table S3.** Colocalization analysis of STRs and the NPC for Figure 2E.

| Sequence | Colocalization # | Colocalization % | No colocalization # | No colocalization % | Total number of cells | p-value relative to no repeats |
| --- | --- | --- | --- | --- | --- | --- |
| No repeats | 52 | 25 | 154 | 75 | 206 | - |
| (CAG)<sub>68</sub> | 74 | 38 | 123 | 62 | 197 | 0.0052 |
| (CTG)<sub>63</sub> | 59 | 42 | 80 | 58 | 139 | 6.3 x10<sup>-4</sup> |
| (TGAAA)<sub>82</sub> | 69 | 54 | 59 | 46 | 128 | 1.2 x10<sup>-7</sup> |
| (TTTCA)<sub>81</sub> | 85 | 44 | 110 | 56 | 195 | 7.9 x10<sup>-5</sup> |
| (GAA)<sub>160</sub> | 64 | 41 | 94 | 59 | 158 | 0.0014 |
| (TTC)<sub>120</sub> | 65 | 43 | 86 | 57 | 151 | 3.1 x10<sup>-4</sup> |

**Table S4.** Zoning analysis for Figure 2F.

| Treatment | STR | Phase | Z1 # | Z1 % | Z2 # | Z2 % | Z3 # | Z3 % | Total number of cells | p-value |
| --- | --- | --- | --- | --- | --- | --- | --- | --- | --- | --- |
| No treatment | (GAA)<sub>99</sub> | S/G2 | 68 | 49 | 39 | 28 | 32 | 23 | 139 | - |
| 1,6-hexanediol | (GAA)<sub>99</sub> | S/G2 | 28 | 32 | 29 | 33 | 30 | 34 | 87 | 0.0094 |
| 2,5-hexanediol | (GAA)<sub>99</sub> | S/G2 | 37 | 47 | 23 | 29 | 18 | 23 | 78 | 0.47 |

**Table S5.** Zoning analysis for Figure 3C.

| Mutant | STR | Phase | Z1 # | Z1 % | Z2 # | Z2 % | Z3 # | Z3 % | Total cells | p-value (compared to WT) |
| --- | --- | --- | --- | --- | --- | --- | --- | --- | --- | --- |
| Wild type | No repeats | S/G2 | 49 | 32 | 53 | 35 | 49 | 32 | 151 | - |
| *nup84Δ* | No repeats | S/G2 | 65 | 34 | 65 | 34 | 60 | 32 | 190 | 0.41 |
| *nup60Δ* | No repeats | S/G2 | 48 | 34 | 42 | 30 | 50 | 36 | 140 | 0.42 |
| *nup1ΔCt* | No repeats | S/G2 | 49 | 29 | 60 | 36 | 58 | 35 | 167 | 0.32 |
| Wild type | (CAG)<sub>73</sub> | S/G2 | 77 | 46 | 41 | 24 | 50 | 39 | 168 | - |
| *nup84Δ* | (CAG)<sub>73</sub> | S/G2 | 58 | 34 | 59 | 34 | 56 | 32 | 173 | 0.013 |
| *nup60Δ* | (CAG)<sub>73</sub> | S/G2 | 74 | 46 | 53 | 33 | 33 | 21 | 160 | 0.51 |
| *nup1ΔCt* | (CAG)<sub>73</sub> | S/G2 | 73 | 34 | 85 | 40 | 54 | 25 | 212 | 0.016 |
| Wild type | (CTG)<sub>67</sub> | S/G2 | 62 | 42 | 48 | 34 | 36 | 25 | 146 | - |
| *nup84Δ* | (CTG)<sub>67</sub> | S/G2 | 55 | 33 | 61 | 36 | 53 | 31 | 169 | 0.045 |
| *nup60Δ* | (CTG)<sub>67</sub> | S/G2 | 62 | 40 | 42 | 27 | 51 | 33 | 155 | 0.38 |
| *nup1ΔCt* | (CTG)<sub>67</sub> | S/G2 | 73 | 34 | 75 | 35 | 64 | 30 | 212 | 0.077 |
| Wild type | (TGAAA)<sub>81</sub> | S/G2 | 95 | 42 | 64 | 28 | 67 | 30 | 226 | - |
| *nup84Δ* | (TGAAA)<sub>81</sub> | S/G2 | 53 | 36 | 50 | 34 | 43 | 29 | 146 | 0.16 |
| *nup60Δ* | (TGAAA)<sub>81</sub> | S/G2 | 89 | 43 | 55 | 26 | 64 | 31 | 208 | 0.48 |
| *nup1ΔCt* | (TGAAA)<sub>81</sub> | S/G2 | 50 | 36 | 38 | 28 | 49 | 36 | 137 | 0.18 |
| Wild type | (TTTCA)<sub>81</sub> | S/G2 | 106 | 44 | 63 | 26 | 73 | 30 | 242 | - |
| *nup84Δ* | (TTTCA)<sub>81</sub> | S/G2 | 62 | 37 | 60 | 36 | 46 | 27 | 168 | 0.098 |
| *nup60Δ* | (TTTCA)<sub>81</sub> | S/G2 | 70 | 43 | 59 | 37 | 32 | 20 | 161 | 0.52 |
| *nup1ΔCt* | (TTTCA)<sub>81</sub> | S/G2 | 54 | 37 | 46 | 32 | 45 | 31 | 145 | 0.12 |
| Wild type | (GAA)<sub>99</sub> | S/G2 | 68 | 49 | 39 | 28 | 32 | 23 | 139 | - |
| *nup84Δ* | (GAA)<sub>99</sub> | S/G2 | 27 | 36 | 31 | 41 | 17 | 23 | 75 | 0.047 |
| *nup60Δ* | (GAA)<sub>99</sub> | S/G2 | 70 | 53 | 36 | 27 | 26 | 20 | 132 | 0.29 |
| *nup1ΔCt* | (GAA)<sub>99</sub> | S/G2 | 56 | 37 | 40 | 26 | 57 | 37 | 153 | 0.022 |

Wild-type values are same as in Table S2.

**Table S6.** GCR rates for Figure 5B.

| Genotype | Sequence | GCR rate | Standard error | Fold change relative to wild type |
| --- | --- | --- | --- | --- |
| Wild type | No repeats | $1.6 \times 10^{-9}$ | $3.2 \times 10^{-10}$ | 1 |
| *mlp1Δ* | No repeats | $2.0 \times 10^{-9}$ | $4.0 \times 10^{-11}$ | 1 |
| *mlp2Δ* | No repeats | $1.5 \times 10^{-9}$ | $2.2 \times 10^{-10}$ | 1 |
| *mlp1Δ mlp2Δ* | No repeats | $2.2 \times 10^{-8}$ | $5.5 \times 10^{-9}$ | 14 |
| *nup1ΔCt* | No repeats | $6.4 \times 10^{-9}$ | $2.6 \times 10^{-9}$ | 4 |
| *nup2Δ* | No repeats | $1.6 \times 10^{-9}$ | $5.0 \times 10^{-10}$ | 1 |
| *nup60Δ* | No repeats | $8.3 \times 10^{-8}$ | $1.3 \times 10^{-8}$ | 50 |
| *nup84Δ* | No repeats | $6.8 \times 10^{-8}$ | $2.2 \times 10^{-8}$ | 41 |

**Table S7.**
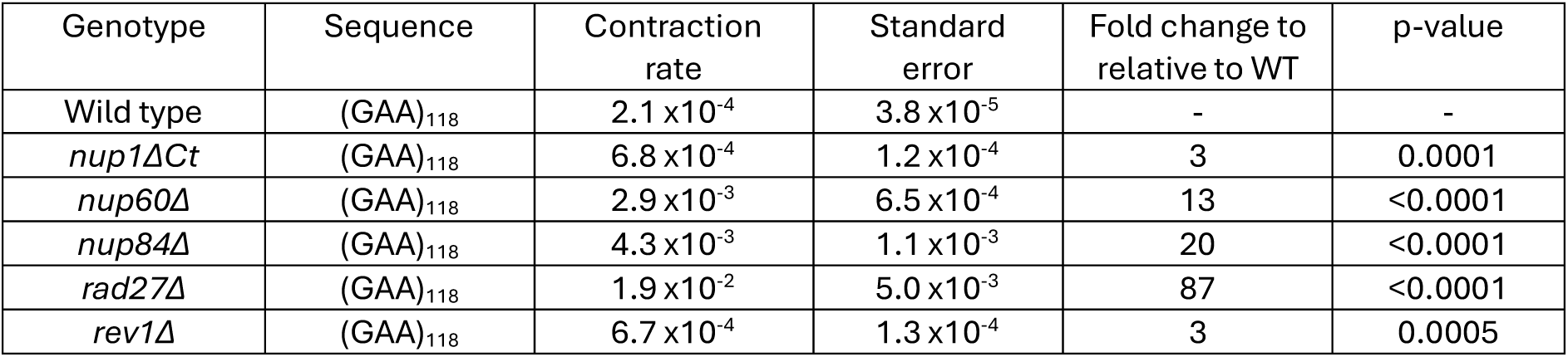
(GAA)_118_ contraction rates for Figure 5C.

| Genotype | Sequence | Contraction rate | Standard error | Fold change to relative to WT | p-value |
| --- | --- | --- | --- | --- | --- |
| Wild type | (GAA)<sub>118</sub> | $2.1 \times 10^{-4}$ | $3.8 \times 10^{-5}$ | - | - |
| *nup1ΔCt* | (GAA)<sub>118</sub> | $6.8 \times 10^{-4}$ | $1.2 \times 10^{-4}$ | 3 | 0.0001 |
| *nup60Δ* | (GAA)<sub>118</sub> | $2.9 \times 10^{-3}$ | $6.5 \times 10^{-4}$ | 13 | <0.0001 |
| *nup84Δ* | (GAA)<sub>118</sub> | $4.3 \times 10^{-3}$ | $1.1 \times 10^{-3}$ | 20 | <0.0001 |
| *rad27Δ* | (GAA)<sub>118</sub> | $1.9 \times 10^{-2}$ | $5.0 \times 10^{-3}$ | 87 | <0.0001 |
| *rev1Δ* | (GAA)<sub>118</sub> | $6.7 \times 10^{-4}$ | $1.3 \times 10^{-4}$ | 3 | 0.0005 |

**Table S8.** (GAA)_100_ expansion rates for Figure 5D.

| Genotype | Sequence | Expansion rate | Standard error | Fold change relative to WT | p-value |
| --- | --- | --- | --- | --- | --- |
| Wild type | (GAA)<sub>100</sub> | $2.1 \times 10^{-5}$ | $7.4 \times 10^{-6}$ | - | - |
| *nup1ΔCt* | (GAA)<sub>100</sub> | $2.1 \times 10^{-5}$ | $6.9 \times 10^{-6}$ | 1 | 0.88 |
| *nup60Δ* | (GAA)<sub>100</sub> | $1.7 \times 10^{-5}$ | $9.8 \times 10^{-6}$ | 0.8 | 0.70 |
| *nup84Δ* | (GAA)<sub>100</sub> | $5.01 \times 10^{-6}$ | $1.0 \times 10^{-6}$ | 0.2 | 0.0005 |
| *rad27Δ* | (GAA)<sub>100</sub> | $1.7 \times 10^{-3}$ | $2.2 \times 10^{-4}$ | 83 | 0.0001 |
| *tof1Δ* | (GAA)<sub>100</sub> | $1.0 \times 10^{-4}$ | $2.2 \times 10^{-5}$ | 5 | 0.0001 |

**Table S9.**
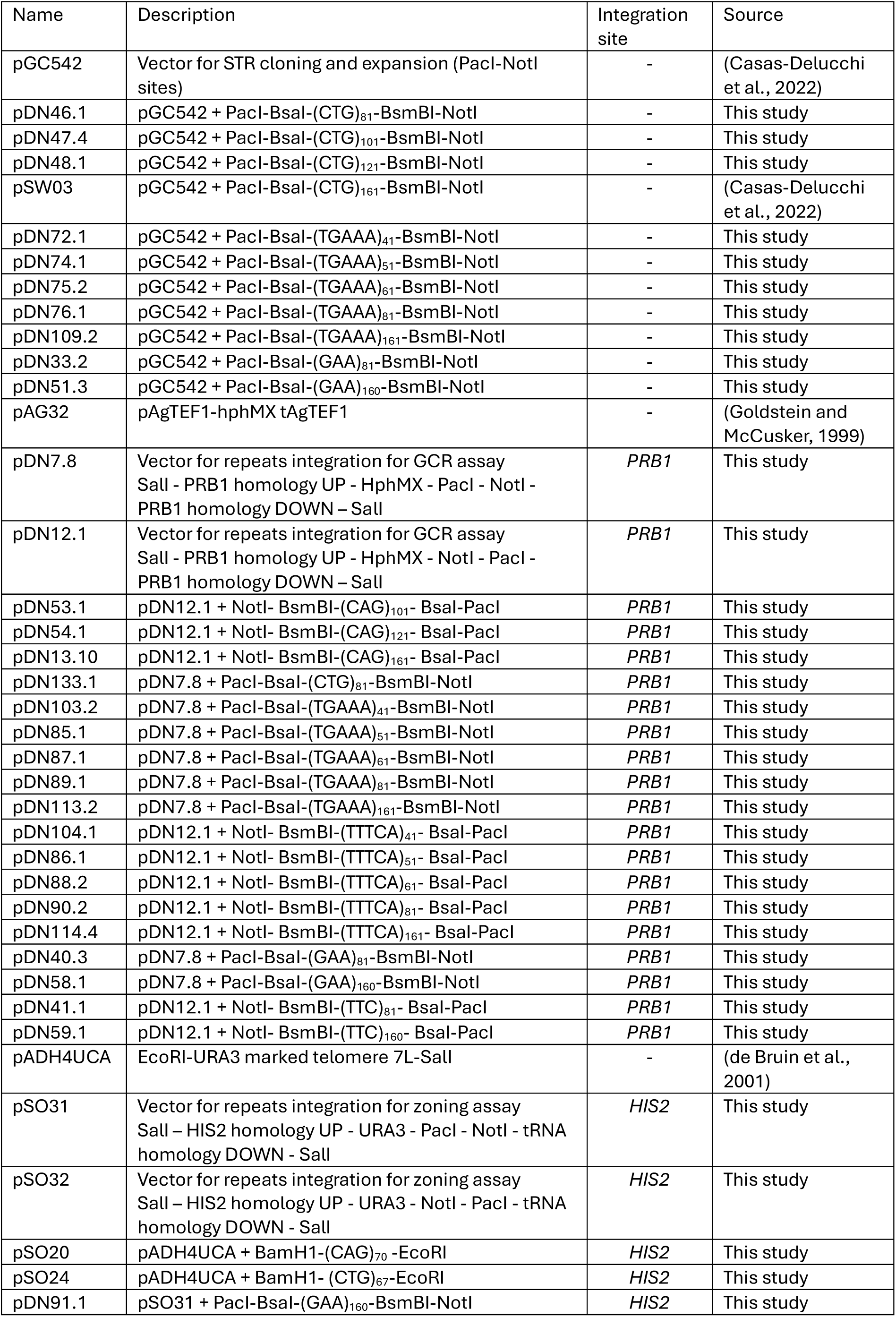
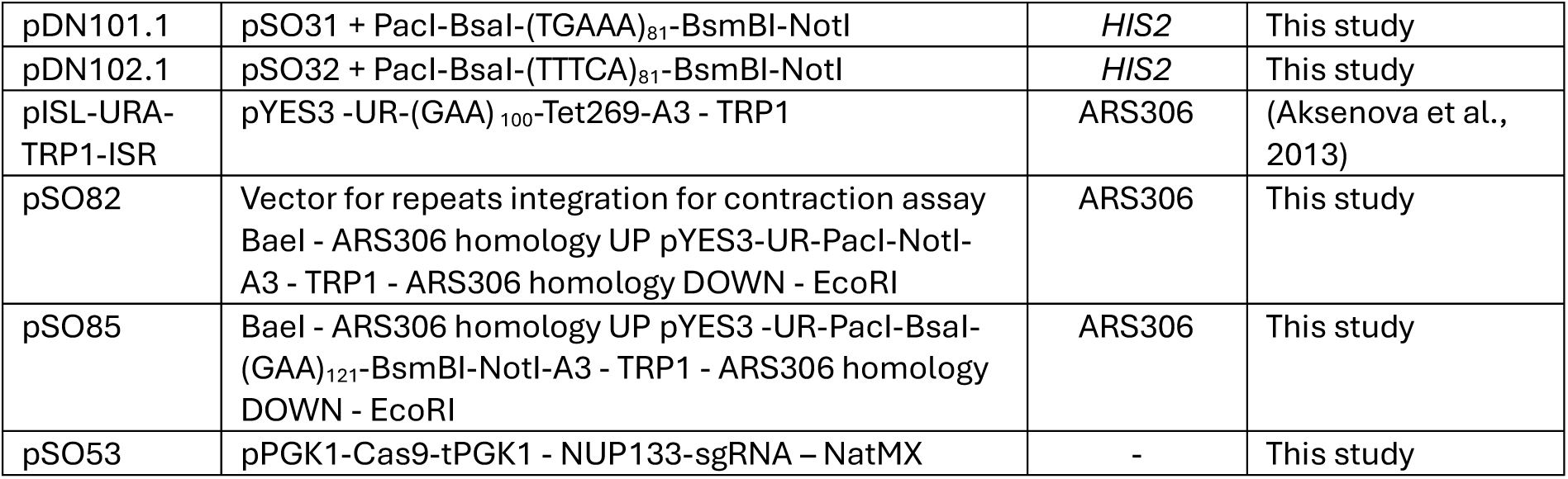
Plasmids used in this study.

**Table S10.**
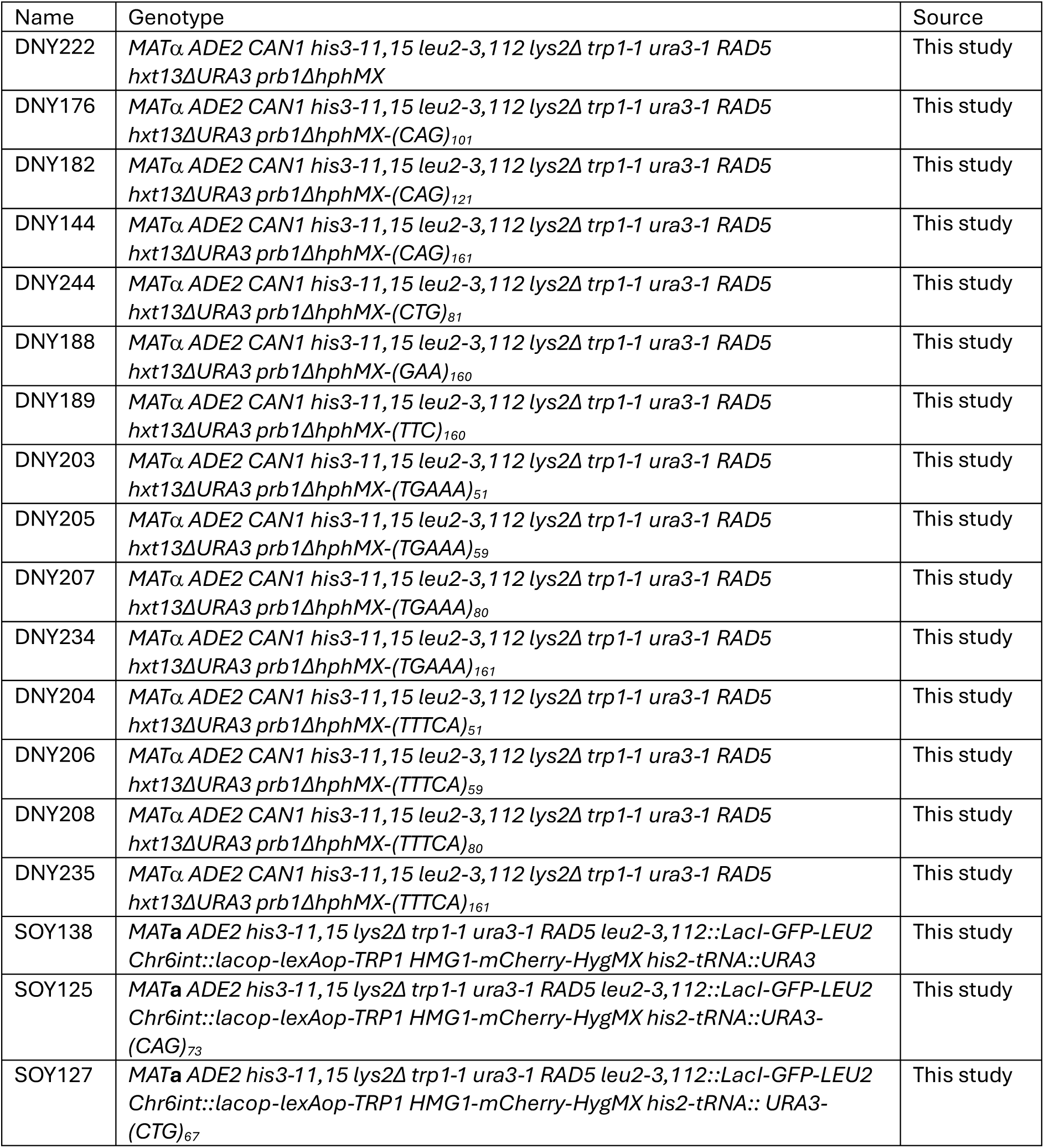
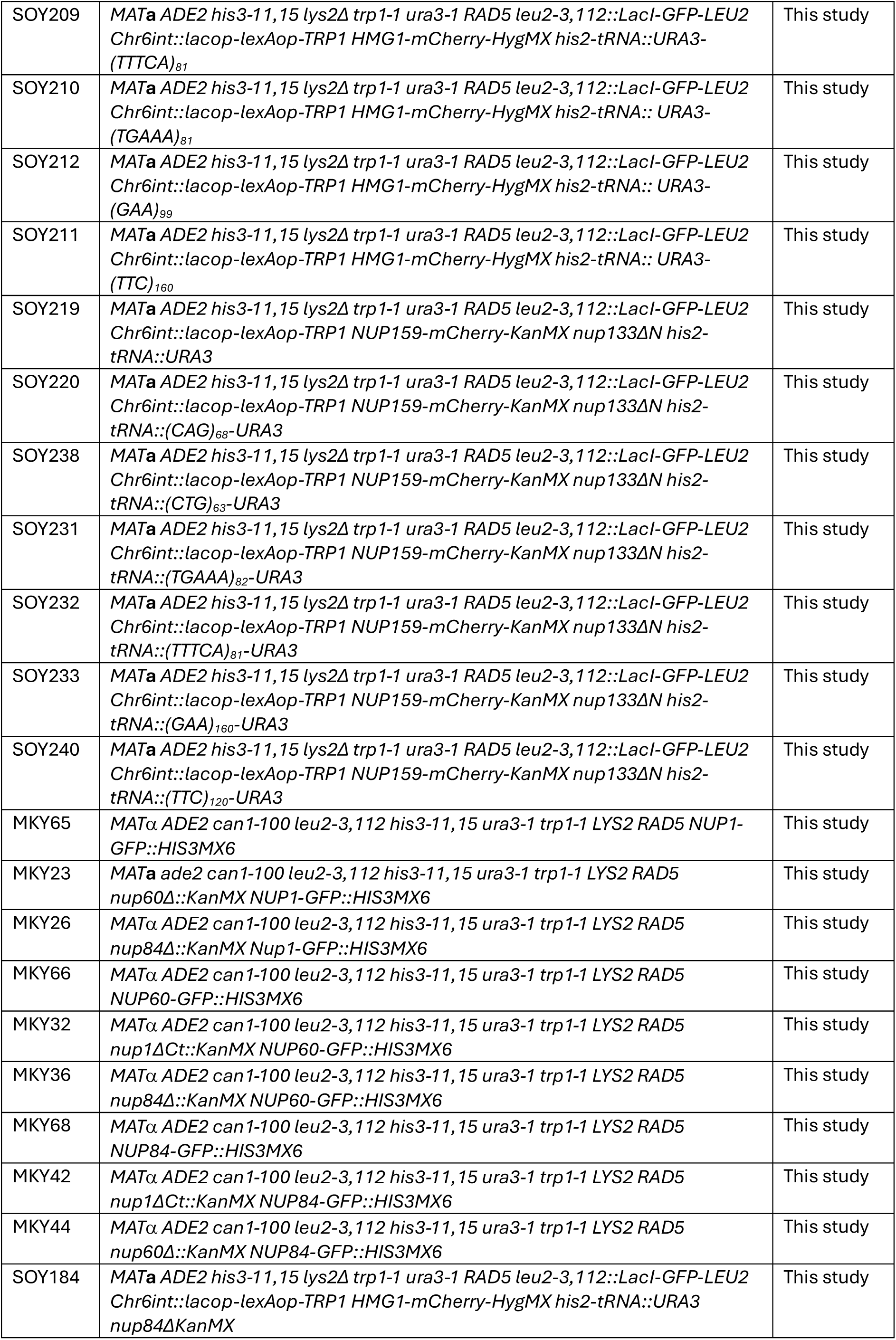
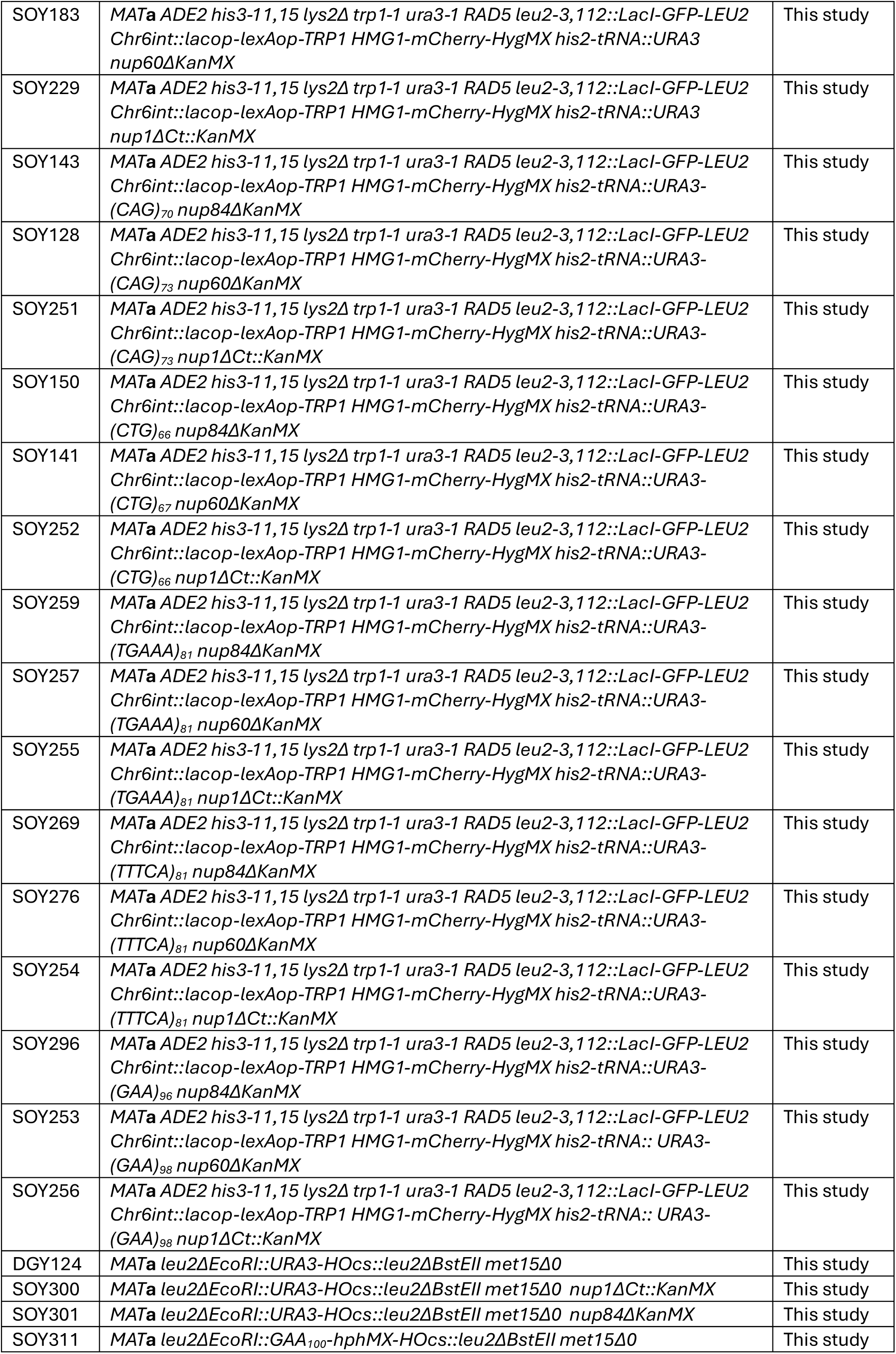
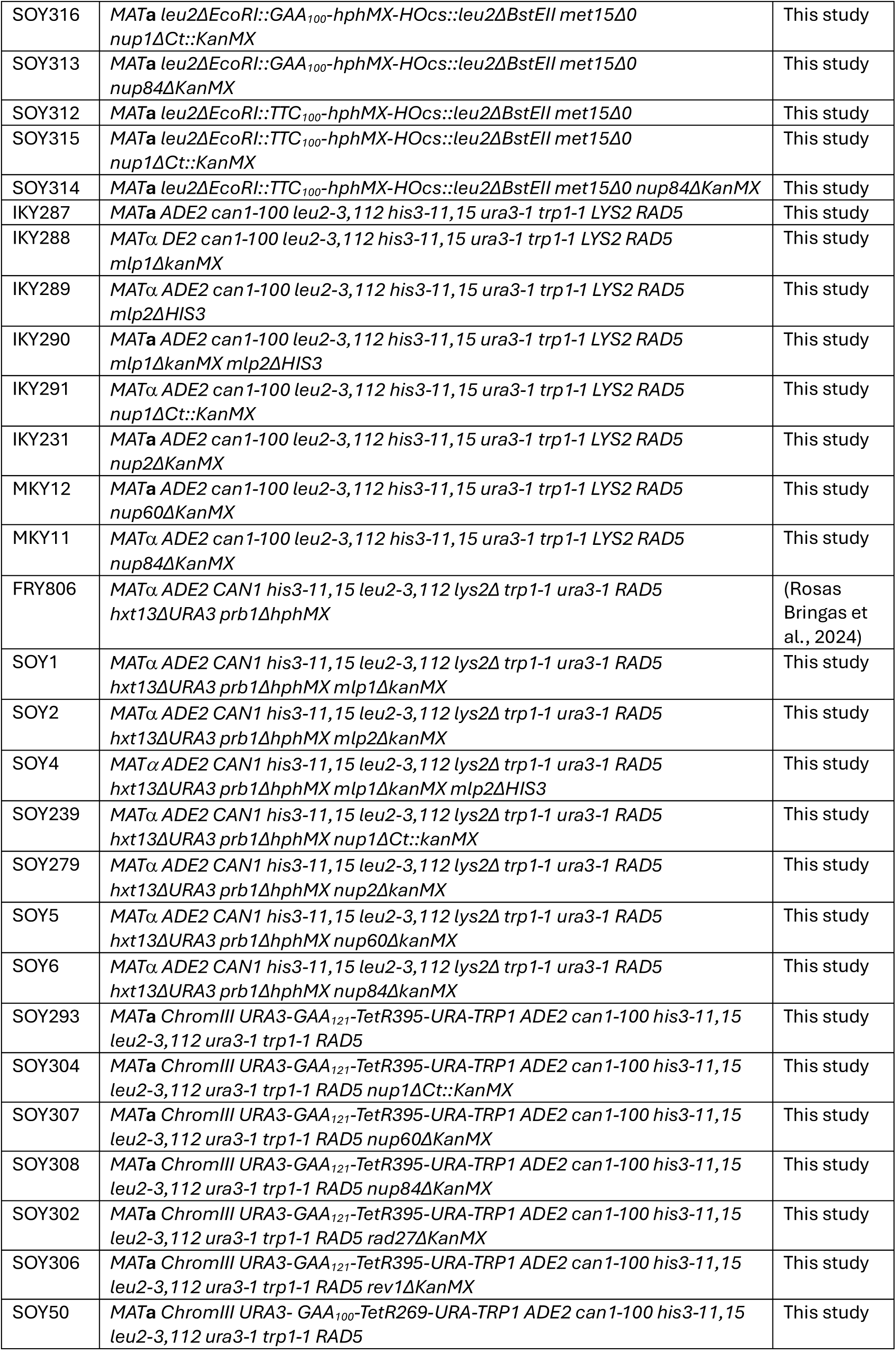
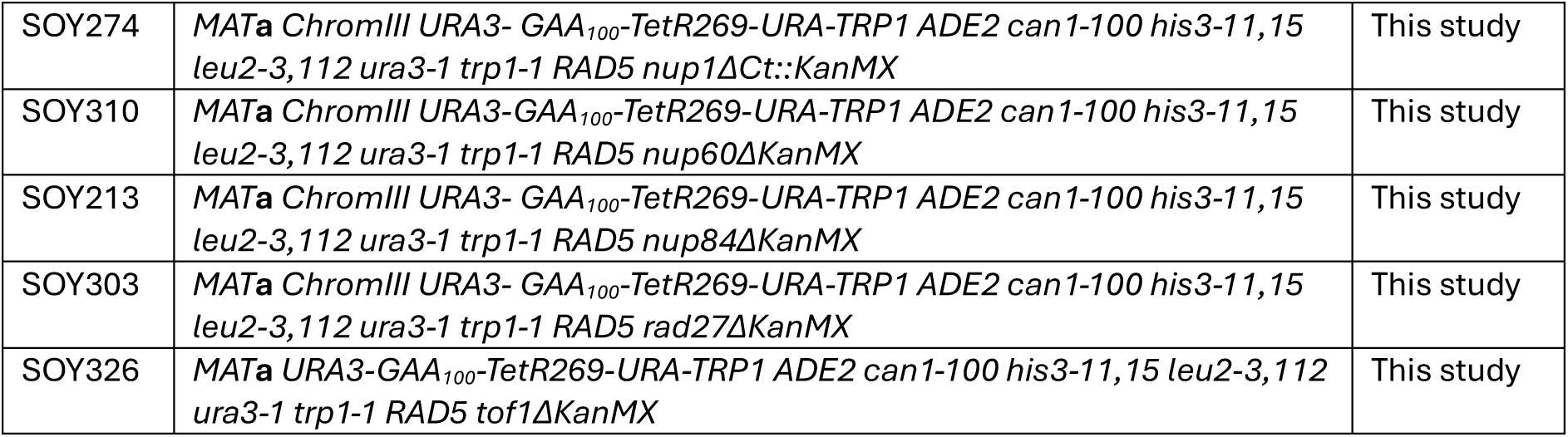
Yeast strains used in this study.

## References

Aguilera, P., M. Dubarry, J. Hardy, M. Lisby, M.N. Simon, and V. Géli. 2022. Telomeric C-circles localize at nuclear pore complexes in *Saccharomyces cerevisiae*. EMBO J. 41:e108736.

Aguilera, P., J. Whalen, C. Minguet, D. Churikov, C. Freudenreich, M.N. Simon, and V. Géli. 2020. The nuclear pore complex prevents sister chromatid recombination during replicative senescence. Nat Commun. 11:160.

Albert, S., M. Schaffer, F. Beck, S. Mosalaganti, S. Asano, H.F. Thomas, J.M. Plitzko, M. Beck, W. Baumeister, and B.D. Engel. 2017. Proteasomes tether to two distinct sites at the nuclear pore complex. Proc Natl Acad Sci U S A. 114:13726–13731.

Alvaro, D., M. Lisby, and R. Rothstein. 2007. Genome-wide analysis of Rad52 foci reveals diverse mechanisms impacting recombination. PLoS Genet. 3:e228.

Beck, M., and E. Hurt. 2017. The nuclear pore complex: understanding its function through structural insight. Nat Rev Mol Cell Biol. 18:73–89.

Bennett, C.B., L.K. Lewis, G. Karthikeyan, K.S. Lobachev, Y.H. Jin, J.F. Sterling, J.R. Snipe, and M.A. Resnick. 2001. Genes required for ionizing radiation resistance in yeast. Nat Genet. 29:426–434.

Brown, R.E., and C.H. Freudenreich. 2021. Structure-forming repeats and their impact on genome stability. Curr Opin Genet Dev. 67:41–51.

Casas-Delucchi, C.S., M. Daza-Martin, S.L. Williams, and G. Coster. 2022. The mechanism of replication stalling and recovery within repetitive DNA. Nat Commun. 13:3953.

Chang, M., M. Bellaoui, C. Boone, and G.W. Brown. 2002. A genome-wide screen for methyl methanesulfonate-sensitive mutants reveals genes required for S phase progression in the presence of DNA damage. Proc Natl Acad Sci U S A. 99:16934–16939.

Chung, D.K., J.N. Chan, J. Strecker, W. Zhang, S. Ebrahimi-Ardebili, T. Lu, K.J. Abraham, D. Durocher, and K. Mekhail. 2015. Perinuclear tethers license telomeric DSBs for a broad kinesin- and NPC-dependent DNA repair process. Nat Commun. 6:7742.

Churikov, D., F. Charifi, N. Eckert-Boulet, S. Silva, M.N. Simon, M. Lisby, and V. Geli. 2016. SUMO-Dependent Relocalization of Eroded Telomeres to Nuclear Pore Complexes Controls Telomere Recombination. Cell Rep. 15:1242–1253.

Cibulka, J., F. Bisaccia, K. Radisavljevic, R.M. Gudino Carrillo, and A. Kohler. 2022. Assembly principle of a membrane-anchored nuclear pore basket scaffold. Sci Adv. 8:eabl6863.

Davis, L.I., and G.R. Fink. 1990. The *NUP1* gene encodes an essential component of the yeast nuclear pore complex. Cell. 61:965–978.

Doye, V., R. Wepf, and E.C. Hurt. 1994. A novel nuclear pore protein Nup133p with distinct roles in poly(A)+ RNA transport and nuclear pore distribution. EMBO J. 13:6062–6075.

Duheron, V., N. Nilles, S. Pecenko, V. Martinelli, and B. Fahrenkrog. 2017. Localisation of Nup153 and SENP1 to nuclear pore complexes is required for 53BP1-mediated DNA double-strand break repair. J Cell Sci. 130:2306–2316.

Dultz, E., and V. Doye. 2025. Opening the gate: Complexity and modularity of the nuclear pore scaffold and basket. Curr Opin Cell Biol. 92:102461.

Erwin, G.S., G. Gursoy, R. Al-Abri, A. Suriyaprakash, E. Dolzhenko, K. Zhu, C.R. Hoerner, S.M. White, L. Ramirez, A. Vadlakonda, A. Vadlakonda, K. von Kraut, J. Park, C.M. Brannon, D.A. Sumano, R.A. Kirtikar, A.A. Erwin, T.J. Metzner, R.K.C. Yuen, A.C. Fan, J.T. Leppert, M.A. Eberle, M. Gerstein, and M.P. Snyder. 2023. Recurrent repeat expansions in human cancer genomes. Nature. 613:96–102.

Fernandez-Martinez, J., J. Phillips, M.D. Sekedat, R. Diaz-Avalos, J. Velazquez-Muriel, J.D. Franke, R. Williams, D.L. Stokes, B.T. Chait, A. Sali, and M.P. Rout. 2012. Structure-function mapping of a heptameric module in the nuclear pore complex. J Cell Biol. 196:419–434.

Feuerbach, F., V. Galy, E. Trelles-Sticken, M. Fromont-Racine, A. Jacquier, E. Gilson, J.C. Olivo-Marin, H. Scherthan, and U. Nehrbass. 2002. Nuclear architecture and spatial positioning help establish transcriptional states of telomeres in yeast. Nat Cell Biol. 4:214–221.

Freudenreich, C.H., and X.A. Su. 2016. Relocalization of DNA lesions to the nuclear pore complex. FEMS Yeast Res. 16.

Gaillard, H., J.M. Santos-Pereira, and A. Aguilera. 2019. The Nup84 complex coordinates the DNA damage response to warrant genome integrity. Nucleic Acids Res. 47:4054–4067.

Gellon, L., S. Kaushal, J. Cebrian, M. Lahiri, S.M. Mirkin, and C.H. Freudenreich. 2019. Mrc1 and Tof1 prevent fragility and instability at long CAG repeats by their fork stabilizing function. Nucleic Acids Res. 47:794–805.

Gemayel, R., M.D. Vinces, M. Legendre, and K.J. Verstrepen. 2010. Variable tandem repeats accelerate evolution of coding and regulatory sequences. Annu Rev Genet. 44:445–477.

Giaever, G., A.M. Chu, L. Ni, C. Connelly, L. Riles, S. Veronneau, S. Dow, A. Lucau-Danila, K. Anderson, B. Andre, A.P. Arkin, A. Astromoff, M. El-Bakkoury, R. Bangham, R. Benito, S. Brachat, S. Campanaro, M. Curtiss, K. Davis, A. Deutschbauer, K.D. Entian, P. Flaherty, F. Foury, D.J. Garfinkel, M. Gerstein, D. Gotte, U. Guldener, J.H. Hegemann, S. Hempel, Z. Herman, D.F. Jaramillo, D.E. Kelly, S.L. Kelly, P. Kotter, D. LaBonte, D.C. Lamb, N. Lan, H. Liang, H. Liao, L. Liu, C. Luo, M. Lussier, R. Mao, P. Menard, S.L. Ooi, J.L. Revuelta, C.J. Roberts, M. Rose, P. Ross-Macdonald, B. Scherens, G. Schimmack, B. Shafer, D.D. Shoemaker, S. Sookhai-Mahadeo, R.K. Storms, J.N. Strathern, G. Valle, M. Voet, G. Volckaert, C.Y. Wang, T.R. Ward, J. Wilhelmy, E.A. Winzeler, Y. Yang, G. Yen, E. Youngman, K. Yu, H. Bussey, J.D. Boeke, M. Snyder, P. Philippsen, R.W. Davis, and M. Johnston. 2002. Functional profiling of the *Saccharomyces cerevisiae* genome. Nature. 418:387–391.

Gietz, R.D., and R.H. Schiestl. 2007. High-efficiency yeast transformation using the LiAc/SS carrier DNA/PEG method. Nat Protoc. 2:31–34.

Hanway, D., J.K. Chin, G. Xia, G. Oshiro, E.A. Winzeler, and F.E. Romesberg. 2002. Previously uncharacterized genes in the UV- and MMS-induced DNA damage response in yeast. Proc Natl Acad Sci U S A. 99:10605–10610.

Horigome, C., D.E. Bustard, I. Marcomini, N. Delgoshaie, M. Tsai-Pflugfelder, J.A. Cobb, and S.M. Gasser. 2016. PolySUMOylation by Siz2 and Mms21 triggers relocation of DNA breaks to nuclear pores through the Slx5/Slx8 STUbL. Genes Dev. 30:931–945.

Jagot-Lacoussiere, L., A. Faye, H. Bruzzoni-Giovanelli, B.O. Villoutreix, J.C. Rain, and J.L. Poyet. 2015. DNA damage-induced nuclear translocation of Apaf-1 is mediated by nucleoporin Nup107. Cell Cycle. 14:1242–1251.

Khadaroo, B., M.T. Teixeira, P. Luciano, N. Eckert-Boulet, S.M. Germann, M.N. Simon, I. Gallina, P. Abdallah, E. Gilson, V. Geli, and M. Lisby. 2009. The DNA damage response at eroded telomeres and tethering to the nuclear pore complex. Nat Cell Biol. 11:980–987.

Khristich, A.N., J.F. Armenia, R.M. Matera, A.A. Kolchinski, and S.M. Mirkin. 2020. Large-scale contractions of Friedreich’s ataxia GAA repeats in yeast occur during DNA replication due to their triplex-forming ability. Proc Natl Acad Sci U S A. 117:1628–1637.

Khristich, A.N., and S.M. Mirkin. 2020. On the wrong DNA track: Molecular mechanisms of repeat-mediated genome instability. J Biol Chem. 295:4134–4170.

Kim, J.C., S.T. Harris, T. Dinter, K.A. Shah, and S.M. Mirkin. 2017. The role of break-induced replication in large-scale expansions of (CAG)*n*/(CTG)*n* repeats. Nat Struct Mol Biol. 24:55–60.

Kölling, R., T. Nguyen, E.Y. Chen, and D. Botstein. 1993. A new yeast gene with a myosin-like heptad repeat structure. Mol Gen Genet. 237:359–369.

Kosar, M., M. Giannattasio, D. Piccini, A. Maya-Mendoza, F. García-Benítez, J. Bartkova, S.I. Barroso, H. Gaillard, E. Martini, U. Restuccia, M.A. Ramirez-Otero, M. Garre, E. Verga, M. Andújar-Sánchez, S. Maynard, Z. Hodny, V. Costanzo, A. Kumar, A. Bachi, A. Aguilera, J. Bartek, and M. Foiani. 2021. The human nucleoporin Tpr protects cells from RNA-mediated replication stress. Nat Commun. 12:3937.

Lea, D.E., and C.A. Coulson. 1949. The distribution of the numbers of mutants in bacterial populations. J Genet. 49:264–285.

Lee, M.E., W.C. DeLoache, B. Cervantes, and J.E. Dueber. 2015. A Highly Characterized Yeast Toolkit for Modular, Multipart Assembly. ACS Synth Biol. 4:975–986.

Lemaître, C., B. Fischer, A. Kalousi, A.S. Hoffbeck, J. Guirouilh-Barbat, O.D. Shahar, D. Genet, M. Goldberg, P. Betrand, B. Lopez, L. Brino, and E. Soutoglou. 2012. The nucleoporin 153, a novel factor in double-strand break repair and DNA damage response. Oncogene. 31:4803–4809.

Liu, L., R.S. Lee, J.M. Twarowski, T. Emagbetere, J. Thomas, J.M. Wells, G.J. Seuferer, K. Lobachev, and A. Malkova. 2025. Genome-wide screen reveals dependence of break induced replication on several distinct checkpoints. Nat Commun.

Liu, S.M., and M. Stewart. 2005. Structural basis for the high-affinity binding of nucleoporin Nup1p to the *Saccharomyces cerevisiae* importin-beta homologue, Kap95p. J Mol Biol. 349:515–525.

Loeillet, S., B. Palancade, M. Cartron, A. Thierry, G.F. Richard, B. Dujon, V. Doye, and A. Nicolas. 2005. Genetic network interactions among replication, repair and nuclear pore deficiencies in yeast. DNA repair. 4:459–468.

Maclay, T.M., J.M. Whalen, M.J. Johnson, and C.H. Freudenreich. 2025. The DNA replication checkpoint targets the kinetochore to reposition DNA structure-induced replication damage to the nuclear periphery. Cell Rep. 44:116083.

Matsuoka, S., B.A. Ballif, A. Smogorzewska, E.R. McDonald, 3rd, K.E. Hurov, J. Luo, C.E. Bakalarski, Z. Zhao, N. Solimini, Y. Lerenthal, Y. Shiloh, S.P. Gygi, and S.J. Elledge. 2007. ATM and ATR substrate analysis reveals extensive protein networks responsive to DNA damage. Science. 316:1160–1166.

Meister, P., L.R. Gehlen, E. Varela, V. Kalck, and S.M. Gasser. 2010. Visualizing yeast chromosomes and nuclear architecture. Methods Enzymol. 470:535–567.

Murat, P., G. Guilbaud, and J.E. Sale. 2020. DNA polymerase stalling at structured DNA constrains the expansion of short tandem repeats. Genome Biol. 21:209.

Nagai, S., K. Dubrana, M. Tsai-Pflugfelder, M.B. Davidson, T.M. Roberts, G.W. Brown, E. Varela, F. Hediger, S.M. Gasser, and N.J. Krogan. 2008. Functional targeting of DNA damage to a nuclear pore-associated SUMO-dependent ubiquitin ligase. Science. 322:597–602.

Neil, A.J., M.U. Liang, A.N. Khristich, K.A. Shah, and S.M. Mirkin. 2018. RNA-DNA hybrids promote the expansion of Friedreich’s ataxia (GAA)n repeats via break-induced replication. Nucleic Acids Res. 46:3487–3497.

Niepel, M., K.R. Molloy, R. Williams, J.C. Farr, A.C. Meinema, N. Vecchietti, I.M. Cristea, B.T. Chait, M.P. Rout, and C. Strambio-De-Castillia. 2013. The nuclear basket proteins Mlp1p and Mlp2p are part of a dynamic interactome including Esc1p and the proteasome. Mol Biol Cell. 24:3920–3938.

Niño, C.A., D. Guet, A. Gay, S. Brutus, F. Jourquin, S. Mendiratta, J. Salamero, V. Géli, and C. Dargemont. 2016. Posttranslational marks control architectural and functional plasticity of the nuclear pore complex basket. J Cell Biol. 212:167–180.

Novarina, D., R. Desai, J.A. Vaisica, J. Ou, M. Bellaoui, G.W. Brown, and M. Chang. 2020. A Genome-Wide Screen for Genes Affecting Spontaneous Direct-Repeat Recombination in *Saccharomyces cerevisiae*. G3 (Bethesda).

Novarina, D., A. Koutsoumpa, and A. Milias-Argeitis. 2022. A user-friendly and streamlined protocol for CRISPR/Cas9 genome editing in budding yeast. STAR Protoc. 3:101358.

Oshidari, R., J. Strecker, D.K.C. Chung, K.J. Abraham, J.N.Y. Chan, C.J. Damaren, and K. Mekhail. 2018. Nuclear microtubule filaments mediate non-linear directional motion of chromatin and promote DNA repair. Nat Commun. 9:2567.

Palancade, B., X. Liu, M. Garcia-Rubio, A. Aguilera, X. Zhao, and V. Doye. 2007. Nucleoporins prevent DNA damage accumulation by modulating Ulp1-dependent sumoylation processes. Mol Biol Cell. 18:2912–2923.

Palancade, B., M. Zuccolo, S. Loeillet, A. Nicolas, and V. Doye. 2005. Pml39, a novel protein of the nuclear periphery required for nuclear retention of improper messenger ribonucleoparticles. Mol Biol Cell. 16:5258–5268.

Penzo, A., M. Dubarry, C. Brocas, M. Zheng, R.M. Mangione, M. Rougemaille, C. Goncalves, O. Lautier, D. Libri, M.N. Simon, V. Géli, K. Dubrana, and B. Palancade. 2023. A R-loop sensing pathway mediates the relocation of transcribed genes to nuclear pore complexes. Nat Commun. 14:5606.

Putnam, C.D., S.R. Allen-Soltero, S.L. Martinez, J.E. Chan, T.K. Hayes, and R.D. Kolodner. 2012. Bioinformatic identification of genes suppressing genome instability. Proc Natl Acad Sci U S A. 109:E3251–3259.

Putnam, C.D., and R.D. Kolodner. 2010. Determination of gross chromosomal rearrangement rates. Cold Spring Harb Protoc. 2010:pdb.prot5492.

Pyhtila, B., and M. Rexach. 2003. A gradient of affinity for the karyopherin Kap95p along the yeast nuclear pore complex. J Biol Chem. 278:42699–42709.

Radchenko, E.A., R.J. McGinty, A.Y. Aksenova, A.J. Neil, and S.M. Mirkin. 2018. Quantitative Analysis of the Rates for Repeat-Mediated Genome Instability in a Yeast Experimental System. Methods Mol Biol. 1672:421–438.

Rajan-Babu, I.S., E. Dolzhenko, M.A. Eberle, and J.M. Friedman. 2024. Sequence composition changes in short tandem repeats: heterogeneity, detection, mechanisms and clinical implications. Nat Rev Genet. 25:476–499.

Riquelme Barrientos, E.C., T.A. Otto, S.N. Mouton, A. Steen, and L.M. Veenhoff. 2023. A survey of the specificity and mechanism of 1,6 hexanediol-induced disruption of nuclear transport. Nucleus. 14:2240139.

Shortt, J.A., R.P. Ruggiero, C. Cox, A.C. Wacholder, and D.D. Pollock. 2020. Finding and extending ancient simple sequence repeat-derived regions in the human genome. Mob DNA. 11:11.

Shulga, N., and D.S. Goldfarb. 2003. Binding dynamics of structural nucleoporins govern nuclear pore complex permeability and may mediate channel gating. Mol Cell Biol. 23:534–542.

Simon, M.N., K. Dubrana, and B. Palancade. 2024. On the edge: how nuclear pore complexes rule genome stability. Curr Opin Genet Dev. 84:102150.

Smith, J., and R. Rothstein. 1999. An allele of *RFA1* suppresses *RAD52*-dependent double-strand break repair in *Saccharomyces cerevisiae*. Genetics. 151:447–458.

Strambio-De-Castillia, C., M. Niepel, and M.P. Rout. 2010. The nuclear pore complex: bridging nuclear transport and gene regulation. Nat Rev Mol Cell Biol. 11:490–501.

Su, X.A., V. Dion, S.M. Gasser, and C.H. Freudenreich. 2015. Regulation of recombination at yeast nuclear pores controls repair and triplet repeat stability. Genes Dev. 29:1006–1017.

Tanudisastro, H.A., I.W. Deveson, H. Dashnow, and D.G. MacArthur. 2024. Sequencing and characterizing short tandem repeats in the human genome. Nat Rev Genet. 25:460–475.

Whalen, J.M., N. Dhingra, L. Wei, X. Zhao, and C.H. Freudenreich. 2020. Relocation of Collapsed Forks to the Nuclear Pore Complex Depends on Sumoylation of DNA Repair Proteins and Permits Rad51 Association. Cell Rep. 31:107635.

Williams, S.L., and G. Coster. 2024. Cloning and expansion of repetitive DNA sequences. Methods Cell Biol. 182:167–185.

Wu, S., K. Chen, T. Xu, K. Ma, L. Gao, C. Fu, W. Zhang, C. Jing, C. Ren, M. Deng, Y. Chen, Y. Zhou, W. Pan, and X. Jia. 2021. Tpr Deficiency Disrupts Erythroid Maturation With Impaired Chromatin Condensation in Zebrafish Embryogenesis. Front Cell Dev Biol. 9:709923.

Yao, Y., Z. Yin, F.R. Rosas Bringas, J. Boudeman, D. Novarina, and M. Chang. 2024. Revisiting the role of the spindle assembly checkpoint in the formation of gross chromosomal rearrangements in *Saccharomyces cerevisiae*. Genetics.

Zhang, Y., A.A. Shishkin, Y. Nishida, D. Marcinkowski-Desmond, N. Saini, K.V. Volkov, S.M. Mirkin, and K.S. Lobachev. 2012. Genome-wide screen identifies pathways that govern GAA/TTC repeat fragility and expansions in dividing and nondividing yeast cells. Mol Cell. 48:254–265.

Zhao, X., C.Y. Wu, and G. Blobel. 2004. Mlp-dependent anchorage and stabilization of a desumoylating enzyme is required to prevent clonal lethality. J Cell Biol. 167:605–611.

Aksenova, A.Y., P.W. Greenwell, M. Dominska, A.A. Shishkin, J.C. Kim, T.D. Petes, and S.M. Mirkin. 2013. Genome rearrangements caused by interstitial telomeric sequences in yeast. Proc Natl Acad Sci U S A. 110:19866–19871.

de Bruin, D., Z. Zaman, R.A. Liberatore, and M. Ptashne. 2001. Telomere looping permits gene activation by a downstream UAS in yeast. Nature. 409:109–113.

Goldstein, A.L., and J.H. McCusker. 1999. Three new dominant drug resistance cassettes for gene disruption in Saccharomyces cerevisiae. Yeast. 15:1541–1553.

Rosas Bringas, F.R., Z. Yin, Y. Yao, J. Boudeman, S. Ollivaud, and M. Chang. 2024. Interstitial telomeric sequences promote gross chromosomal rearrangement via multiple mechanisms. Proc Natl Acad Sci U S A. 121:e2407314121.

